# Distinct hippocampal codes emerge during brain-machine interface navigation

**DOI:** 10.64898/2026.05.11.724143

**Authors:** Charles Micou, Hinze Ho, Timothy O’Leary, Julija Krupic

## Abstract

The hippocampus integrates external cues and self-motion to construct cognitive maps. Activating these maps independently of immediate sensory and motor signals can support navigation by predicting future locations. The neural basis of such internally driven activation remains poorly understood. To address this, we had mice use a brain-machine interface (BMI) to directly control navigation from real-time hippocampal activity. In this condition, CA1 responses encoding running movement were not navigationally relevant, and place codes rapidly reconfigured to form new representations that discarded locomotion-related signals. By comparing neural representations across BMI-controlled navigation, locomotion-controlled navigation, and passive playback of predetermined routes, we found evidence of response patterns that were specific to conditions in which neural activity could causally influence an animal’s travel. This suggests the existence of CA1 responses that are not tied to external stimuli or self-motion, and that are suppressed when animals are merely passive observers.

## Main

The hippocampus is proposed to act as a cognitive mapping system that integrates information about locations and sensory experiences, as well as events and executed actions associated with those locations^1,2^. Place cells have been identified as fundamental units of this map and are postulated to be activated in one of two ways: from exteroceptive sensory cues or from motor-related signals corresponding to locomotion^3^. These ideas enjoy wide experimental support^4–9^.

However, the encoding of action plans^10^ and progress toward goals^11,12^ suggest that the hippocampus is also essential for predictive computations^13^. Maps that link an animal’s actions to transitions between states are therefore a standard formulation in modern computational models of the hippocampus^14–16^. In these frameworks, prediction and evaluation of future states would require driving hippocampal activity not with immediate sensory stimuli or afferent motor actions, but with internally generated activity that signals the ability and need to make a navigational choice. Given the critical role of the hippocampus in navigation and its sensitivity to task engagement^17–20^, we would expect such signals to manifest most strongly in contexts where animals actively navigate. However, isolating internally generated signals is technically challenging, as navigation actions are typically tightly coupled to motor actions.

To address this question, we developed an experimental paradigm for manipulating the causal relation between hippocampal representations and navigational outcomes, independently of locomotion, by using a Virtual Reality (VR) navigation task coupled to a real-time hippocampal Brain Machine Interface (BMI). This setup allowed us to explicitly manipulate whether hippocampal activity influences navigation, and to manipulate the relation between motor actions and perceived VR motion. We focused specifically on CA1, given both the wealth of existing work exploring responses in this region^21,22^ and its proposed role in identifying discrepancies between predicted and actual outcomes^2,23,24^.

## Results

### A brain-machine interface for internally driven navigation

We developed a VR task in which head-fixed mice navigated through the same environment under two different movement conditions. Initially, navigation was controlled by locomotion (running on a wheel), but this could be switched to control via a BMI that decoded from neural activity in CA1 in real time (Fig. 1a). This offered an experimental means of manipulating the causal relationship between CA1 signals and navigation by producing a task in which performance depended directly on the hippocampus maintaining a consistent cognitive map. Animals traversed a 400 cm, one-dimensional virtual track, rich in both proximal and distal visual cues, with a reward site fixed at 250 cm (Fig. 1b, top). Expert animals anticipated rewards and adopted both anticipatory licking and stereotypical running trajectories between laps of the track, slowing down before stopping at the reward site (Fig. 1b, middle).

**Fig. 1.**
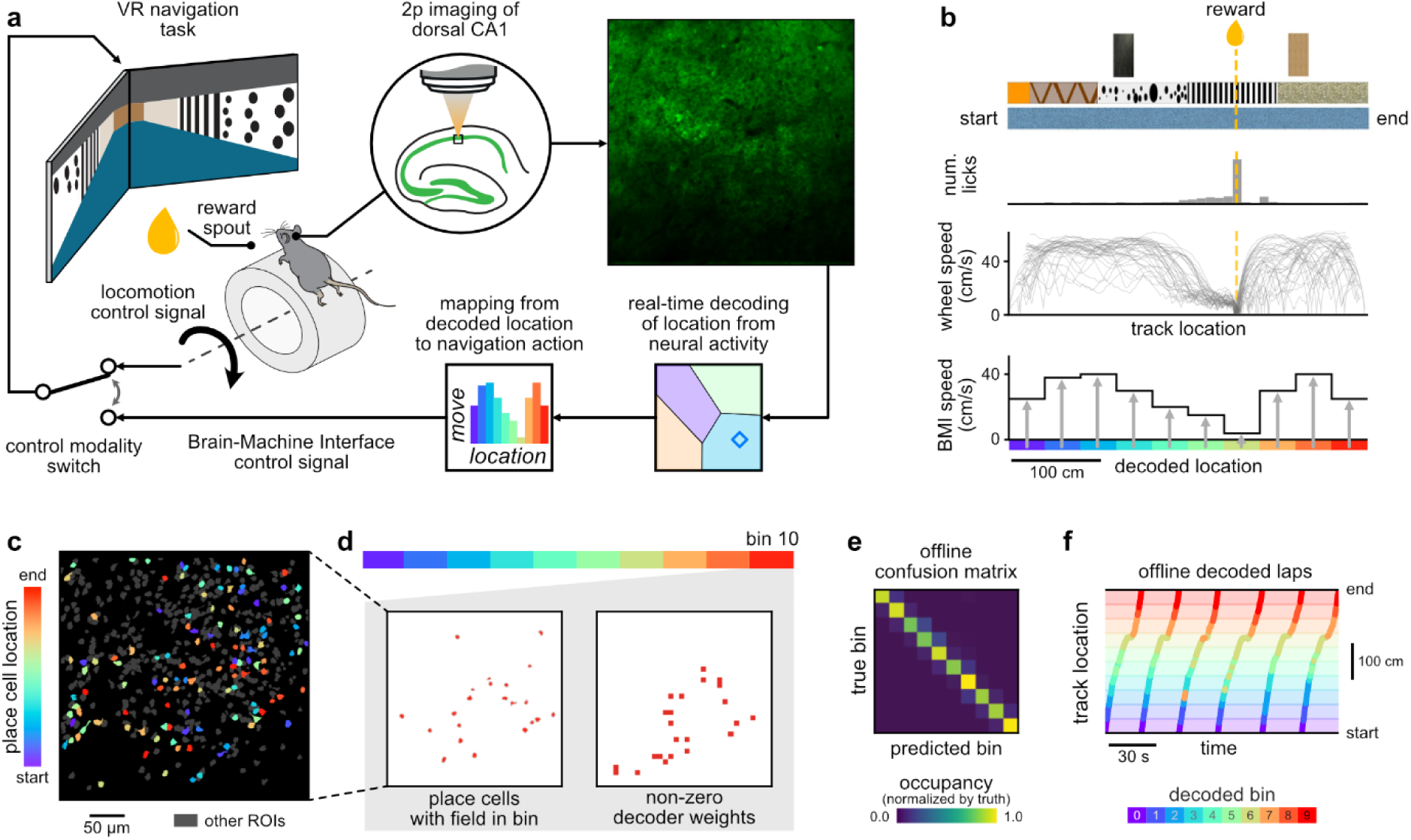
A real-time hippocampal BMI yokes population activity to movement through a VR environment, allowing animals to solve the same navigation task both with and without needing to run to navigate. **(a)** Overview schematic showing two movement conditions that can be switched between. Under locomotion, animals run on the wheel to move through the VR environment. In BMI, the activity of a population of hundreds of cells in dorsal CA1 is imaged at 30 Hz. Each frame is first decoded to extract the location of highest neural similarity to training data and then transformed to movement through the environment. **(b)** The navigation task consists of a 400 cm track with proximal visual cues on the track walls and distally visible landmark towers (top). Animals learn to stop at a fixed reward site, with experienced animals developing anticipatory licking and a stereotypical ‘ballistic’ velocity trajectory (middle). In BMI, a perfectly accurate decoder could reproduce these trajectories by mapping a decoded spatial location to the corresponding stereotypical speed (bottom). **(c)** An example FoV showing a dorsal CA1 population containing place cells. **(d)** We use decoders based on SVMs, which allow inspection of which features are most informative for decoding each class (bin). After training, these correspond closely to place cells active in the same bin. **(e)** An example confusion matrix, normalized by ground truth, showing decoder accuracy for decoding animal position quantized to 10 spatial bins. Decoder trained on 75% of laps from a 10-minute session and evaluated on unseen laps. **(f)** Evaluation of the decoder’s accuracy over the course of six unseen laps of the track.

Our BMI yoked traversal velocity in the VR environment to the activity of hundreds (250-500) of CA1 neurons via two-photon Calcium imaging at 30 Hz (Fig. 1a, Supplementary Fig. 1a). The BMI consisted of three stages. The first stage, signal processing, stabilized the image and extracted changes in Ca^2+^ activity (Supplementary Fig. 1b). The second stage, a decoder, used a support vector machine (SVM)^25^ to predict the animal’s location to within one of ten 40 cm spatial bins directly from a single frame of Ca^2+^ activity (Supplementary Fig. 2). The final BMI stage mapped decoded locations to actions by generating a forward VR movement, subject to a maximum acceleration between frames, chosen as double the maximum acceleration observed in running mice to mitigate sudden, discontinuous motion that could potentially disorient or distract the animal.

The location-to-speed mapping was identical for all mice and was based on empirically measured average running speeds at each track position collected from expert-level performance of the locomotion VR task (Fig. 1b, bottom). By construction, accurate BMI decoding of position at each moment would therefore mimic the motion of expertly executed laps. Furthermore, the SVM decoder weights allowed clear identification of which neurons were critical for decoding each location on the track, and these corresponded closely with place cells (Fig. 1d). We stress that, in contrast to traditional BMI experiments which attempt to explore ideas of volitional control via an interface^26^, the objective of this manipulation is to provide a real-time causal coupling between consistent hippocampal representations and traversals through the environment.

To understand the consequence of poor BMI decoding, we simulated the effect of uniform, random neural activation on BMI decoder accuracy. Due to the relative class volumes of the SVM decoder, this produced a random distribution of output speeds that was biased towards the slowest possible movement through the environment (Supplementary Fig. 2b, Supplementary Text). This outcome would therefore behaviorally incentivize a consistent hippocampal representation during BMI, because more rapid progress outside the reward zone would enable greater reward in the time window of the experiment. Given the high accuracy of the BMI decoder when validated offline on locomotion-based laps of the track (Fig. 1e-f), we expected the decoder to produce consistent, well-executed laps of the track via the BMI, provided that the neural representation of the environment remained stable.

### Retraining decoders on BMI-specific hippocampal codes

We first asked if mice could successfully perform the navigation task via BMI control. To test this, we used experimental sessions of four 10-minute blocks that followed an ‘ABBA’ structure (Fig. 2a). The VR environment was identical for all blocks. The first and last blocks consisted of baseline blocks in which VR motion was controlled by running on the wheel. During the middle blocks navigation was controlled via the BMI. This design allowed us to assess changes in neural representation of the environment both during and following BMI usage (Fig. 2b).

**Fig. 2.**
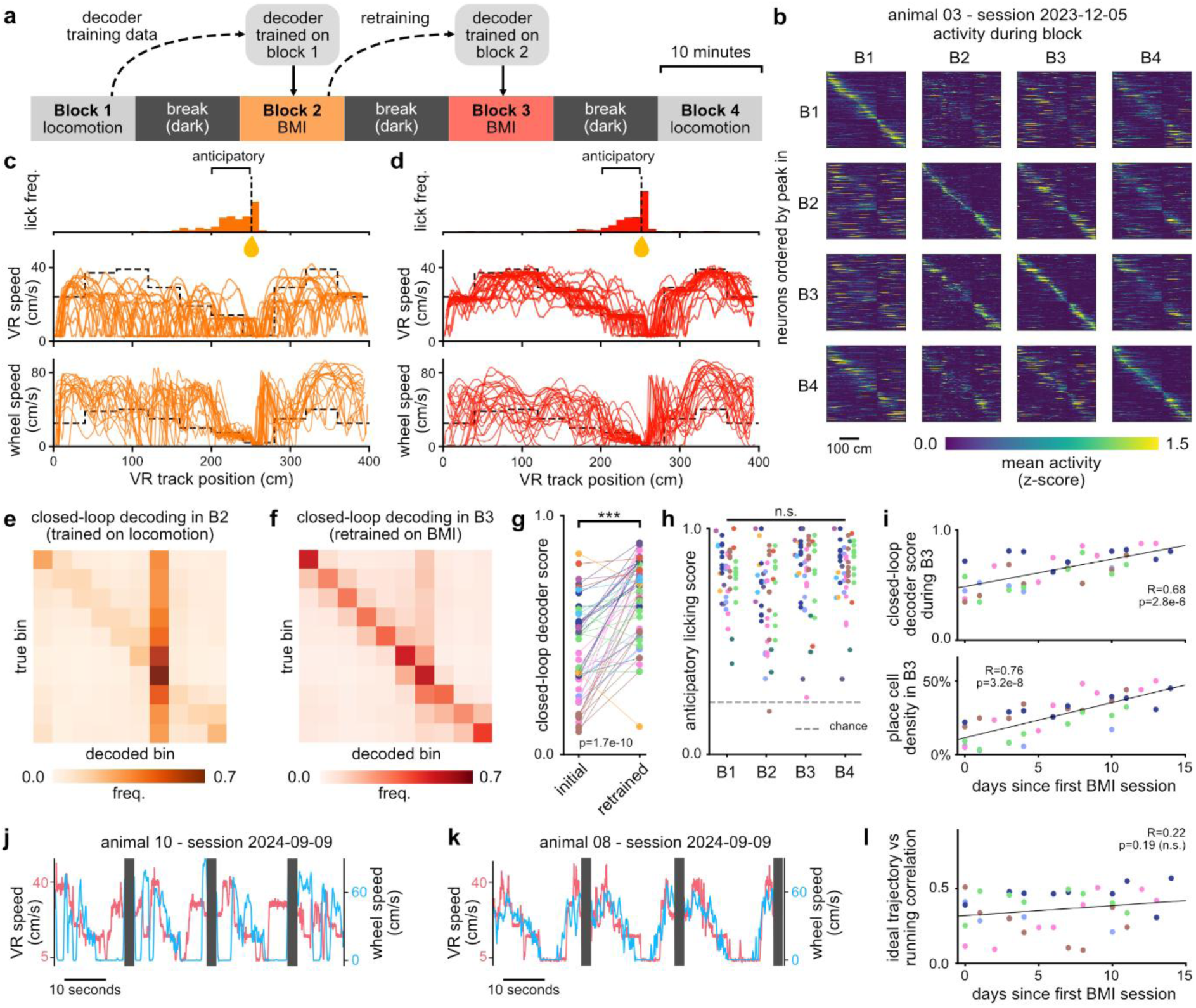
Neural representations of the same task differ when performed via locomotion and via BMI. Good closed-loop BMI performance can be recovered by retraining decoders on these new representations. **(a)** Experimental sessions consist of four 10-minute imaging blocks interleaved with 10 minutes in darkness. Each BMI block uses the previous block as training data for its decoder. **(b)** Example population ratemaps for a single session, showing the mean dF/F activity of 352 neurons at each location on the track. **(c)** Behavior during block 2 of the session in (b), showing the spatial distribution of spout licking (top); the animal’s speed through the VR environment as controlled by BMI, with each lap shown as a trace and the target trajectory indicated by a dashed line (middle); and the speed of wheel manipulation (bottom). **(d)** As in (c), shown for block 3 following retraining of the decoder. **(e)** Confusion matrix illustrating decoder accuracy during block 2, normalized by ground truth and aggregated over all 50 sessions (11 animals). **(f)** As in (e), shown for block 3. **(g)** Comparison of closed-loop decoder accuracy before and after retraining. One point pair per session, color indicates animal identity. **(h)** Anticipatory licking accuracy in each session. **(i)** Decoder score (top) and the fraction of neurons in the population with place fields (bottom) during block 3 for a cohort of five animals repeating the task over two weeks. **(j)** Sample laps for an animal with low correlation between wheel manipulation speed (blue) and BMI-driven speed (red). Darkness interval of 2 seconds between laps shown in gray. **(k)** As in (j), but for an example animal with high correlation between the two. **(l)** For the same cohort as (i), a plot of the correlation between wheel manipulation speed and the idealized trajectory used by the BMI.

We trained an initial BMI decoder using imaged CA1 activity during the first baseline block. Closed-loop performance of the task with BMI was poor, despite high offline decoder accuracy, unchanged sensory cues and the animal’s locomotion being unimpeded (it was still free to run on the wheel, even when this motion was decoupled from VR control). Laps did not adhere to the target trajectory and progressed haphazardly (Fig. 2c). This was explained by poor decoding: inspection of the decoder confusion matrix revealed a failure to decode track location, biasing predictions toward the lowest velocity bin (Fig. 2e), similar the outcome of decoding simulated random activity (Supplementary Fig. 2b). We therefore reasoned that the representation of the same environment differed sufficiently between the locomotion and BMI conditions to collapse decoder accuracy. Based on this hypothesis, we used the poorly-executed BMI laps and corresponding neural data to retrain the decoder between blocks 2 and 3. This ‘second generation’ BMI decoder showed significantly improved performance on the following BMI block and produced well-executed laps (Fig. 2d). Crucially, this improved BMI decoding also enabled our desired condition of directly yoking navigation to hippocampal maps.

Over the course of 50 sessions performed by 11 mice (Supplementary Fig. 3), decoder retraining consistently produced accurate decoders that were improvements over their first-generation counterparts (Fig. 2e-g), suggesting that the neural responses formed in the first BMI block remained stable (Fig. 2b) despite significant changes in the animal’s trajectory and apparent discrepancies between the locomotion signal (how it manipulated the wheel) and VR motion. We used anticipatory licking of the reward spout as an indicator of an animal’s engagement and awareness of its location. We found that animals remained engaged and could self-locate despite the potential for frustration with poorly-executed BMI laps during the initial BMI block (Fig. 2h, Supplementary Fig. 4). Furthermore, we found that the BMI performance of individual animals tended to increase with experience, accompanied by an increasing density of place fields during the BMI blocks in successive sessions (Fig. 2i) despite extensive familiarity with this VR task in the locomotion condition prior to BMI trials (see Methods). The relationship between the number of available place cells and decoder accuracy (Supplementary Fig. 4d) is potentially self-reinforcing, as the more consistent experiences provided by high-accuracy decoders makes it easier to form stable place codes, while the increased density of place cells provides a richer and more redundant feature space from which to decode.

We allowed the mice to move on the wheel in all conditions. This meant that animals had no physical cue to determine how motion was controlled, and allowed us to quantify the disparity between locomotion movement and visual feedback. Because wheel manipulation does not affect movement during BMI, it becomes an unconstrained degree of freedom in the task. Accordingly, some animals exhibited significant discrepancies, stopping on the wheel entirely during segments of VR motion (Fig. 2j). Others, in contrast, continued to run in close agreement with their learned stereotyped trajectory (Fig. 2k). Crucially, animals were able to achieve high BMI performance across the full range of cases, from complete mismatch between wheel manipulation and VR motion through to close correlation between both (Supplementary Fig. 4e). The neural representation itself appears insensitive to wheel manipulation during BMI (Supplementary Fig. 5a-b), suggesting that these representations do not rely on locomotion-based path integration, and any residual correlation between matched speeds and decoder accuracy might instead be attributed to task engagement (Supplementary Fig. 4f-g, Supplementary Text). Finally, unlike place cell density and BMI decoder accuracy, adherence to the target trajectory did not meaningfully vary with experience (Fig. 2l). Taken together, these observations suggest that hippocampal activity can drive BMI motion independently of putative afferent or reafferent locomotion signals.

### BMI-controlled navigation induces rapid partial reconfiguration of CA1 maps

To quantify discrepancies in representation between the locomotion and the BMI conditions, we computed the mean population vector correlation (PVC) between each pair of blocks (Fig. 3a). As expected, we found that the mutual correlation between blocks of the same type of movement was greater than between blocks with different movement modalities (p=7.5e-5, 2-sided t-test corrected for animal identity), pointing to the rapid emergence of a distinct place cell code in the BMI condition. However, and surprisingly, the representational similarity between conditions was not symmetrical with respect to time: the first baseline and BMI blocks were less mutually similar than the final BMI and baseline blocks (B1B2 vs B3B4, p=0.02, pairwise 2-sided t-test corrected for animal identity and accounting for multiple comparisons using a Bonferroni correction). This increase in similarity cannot be explained purely by an increase in the number of place cells during the final baseline block, as the fraction of the population participating in the representation does not increase over the course of the session—in fact, it slightly decreases (Fig. 3b). A more parsimonious explanation is that neuron-level retuning results from exposure to the BMI condition. Inspection of individual dF/F traces revealed particular tuning patterns in certain neurons that might explain the asymmetry in PVC comparisons (Fig. 3c). These neurons formed place fields in two different locations: they exhibited activity at one location during the initial locomotion block, at a distinct and different location during the BMI block, and then at both locations simultaneously upon returning to the locomotion condition. We refer to these tuning curves as ‘superposed’, as their activity in the final baseline appears at the combination of locations from the previous condition. At the population level, this superposition appears consistent throughout the final block, rather than being a transient time-sensitive effect (Supplementary Fig. 5c).

**Fig. 3.**
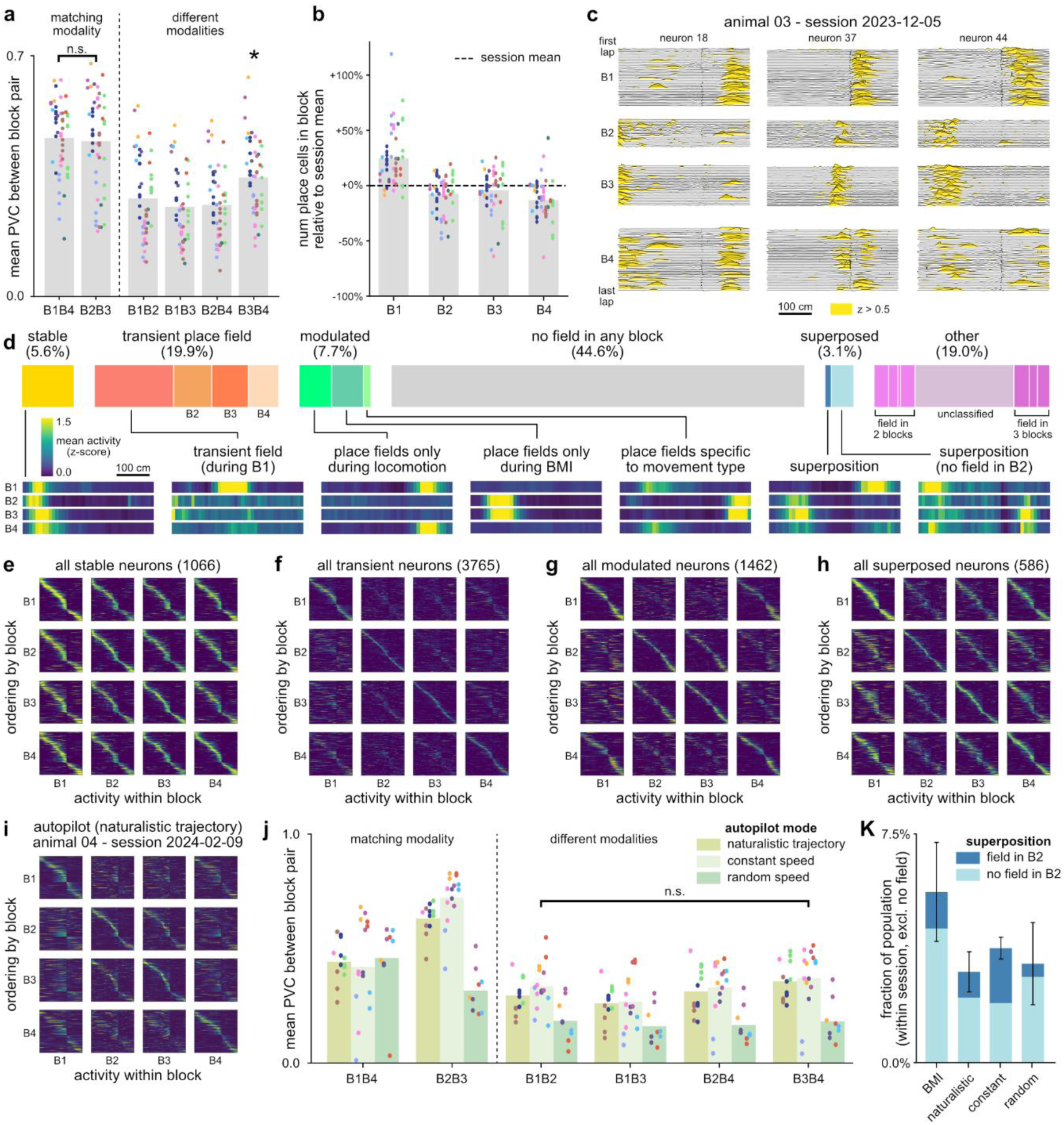
After widespread remapping when entering the BMI condition, CA1 forms new fields that persist after returning to the locomotion condition. These can linearly superpose with the existing cognitive map. **(a)** Similarity of representations between each pair of blocks as measured by mean PVC. Bars show mean over all sessions. **(b)** The number of neurons with a place field, shown for each block in a session relative to the mean number of such neurons across the session. **(c)** dF/F traces for each lap for three example neurons exhibiting superposed fields in block 4. Traces have been low-pass filtered and highlighted during periods of elevated activity for legibility. **(d)** Relative frequencies of heuristic categories of neuron response based on place field locations within each block (18902 neurons, 50 sessions). Neuron ratemaps illustrate archetypal examples for each category. **(e)** Population ratemap, as in Fig. 2b, synthesized from all 50 sessions using only neurons in the stable category. Shown also for **(f)** transient fields, **(g)** fields modulated by the movement condition, and **(h)** fields that linearly superpose in the final block. **(i)** A population ratemap for a control session in which BMI blocks (B2 and B3) were replaced by a passive motion condition that mimics naturalistic lap execution. **(j)** Mean PVCs between block pairs, as in (a), shown for a variety of passive motion conditions including naturalistic trajectories, constant velocity along the track, and random forward velocity. **(k)** Fraction of place field-forming neurons in the population categorized as exhibiting superposition, shown for the BMI and passive motion sessions. Bars indicate mean over sessions, error bars are +/- SEM.

To provide an intuition for how neuron-level tuning might correspond to the population-level effect, we devised a heuristic classification scheme to group neurons according to the conditions in which they formed place fields (Fig. 3d). A small proportion (5.6%, 1066 examples) of all cells (including those that remain silent but can be observed using two-photon imaging, which was on average 44.6% of all ROIs classified as cells) retained stable place fields across baseline and BMI blocks (Fig. 3d-e). This is consistent with our observation that mice continued either to lick in anticipation of reward during BMI usage (Fig. 2d, top) or, in some instances, to manipulate the wheel in accordance with their stereotyped trajectory (Fig. 2k, Supplementary Fig. 4e); pointing to an awareness of their self-location with respect to the reward and environmental cues, and consistent with previous studies demonstrating that place cell activity plays a causal role in an animal’s perception of location^27,28^. Given the uninformative nature of path-integrating locomotion in the BMI condition, we suggest that these cells are primarily driven by visual sensory input, with the proportion of these putative visually driven cells consistent with that previously reported^4^.

Many place cells (19.9%, 3765 examples) were identified only in a single block and were transient in nature (Fig. 3d, f). Neurons that formed stable fields across multiple blocks were often modulated by navigation condition (Fig. 3d, g), either shifting field locations between baseline and BMI blocks (0.8%, 155 examples), or exhibiting fields exclusively in the locomotion condition (3.5%, 661 examples) or during BMI control (3.4%, 646 examples). Locomotion-exclusive fields may correspond to cells whose inputs are dominated by path integration signals, consistent with previous work^4^. Cells specific to each condition covered the entirety of the track (Fig. 3g).

We found that a portion of all neurons (3.1%, 586 examples), comparable to the movement type-specific categories, exhibited superposed responses (Fig. 3d, h).

Despite being a relatively small portion of the overall population, these neurons are responsible for approximately 17% of the discrepancy in PVC that follows BMI usage. The definition we used for superposed responses was conservative (see Methods) and excludes marginal cases. When we relaxed the definition by dropping the requirement for a place field in block 1, the larger proportion of matched neurons (6.1%) accounted for a greater proportion (49%) of the discrepancy in PVC (Supplementary Fig. 5d). However, to avoid confounds between time- and BMI-dependent effects, we restrict all subsequent analysis of superposed neurons to the conservative definition.

### Emergence of place field superposition is BMI-specific

The BMI condition is a complex manipulation: while it successfully yokes VR movement directly to hippocampal activity, it also alters the relationship between running and movement, and potentially lowers the rate at which rewards are received. Previous studies have shown that changes in context or an animal’s engagement^18,20,29^, as well as motion error signals arising from a mismatch between locomotion and sensory cues^4,5^, or indeed the manipulation of the gain between wheel running and VR travel^30^, can induce partial or global remapping of place codes in CA1. We therefore asked whether the representations that emerge during BMI usage are specific to a condition in which CA1 activity is causally related to motion, or, alternatively, whether the representations formed in BMI are equivalent to those formed in passive motion, similar to previous findings exploring mismatches between landmark and locomotion signals.

To address this, we ran additional control experiments that replaced BMI-controlled blocks with an open-loop ‘autopilot’ condition that played back predetermined traversals through the VR environment. Unlike the BMI condition, in which place cell dynamics directly controlled navigation, the autopilot condition is passive: hippocampal activity is inconsequential to task success. However, common to both conditions is the mismatch between locomotion and changes in external cues. We used a variety of autopilot velocity profiles: constant speed, random speeds, or naturalistic trajectories that mimicked the mean velocity profiles of well-executed laps (Supplementary Fig. 6a-f). Both constant speed and naturalistic autopilot conditions provide passive yet predictable motion. Consistent with previous studies^4,5^, across animals and autopilot profiles, we found partial place cell remapping in the autopilot condition (Fig. 3i). However, we found no evidence that experience of the autopilot condition altered the locomotion-based representation of the environment, as measured by comparing mean PVC between block pairs B1B2 and B3B4 (Fig. 3j, Supplementary Table 1), and we found fewer instances of neurons heuristically categorized as superposed (Fig. 3k, Supplementary Fig. 6g). Furthermore, while the velocity distributions experienced in the random speed autopilot condition were by construction identical to those in the poorly executed BMI laps from the previous section (see Methods), the quality of place fields formed was higher in the BMI condition than this autopilot condition (Supplementary Fig. 6h-i).

Next, we further addressed the possibility that, due to discrepancies between speed of travel in the BMI and locomotion conditions, animals were merely perceiving the BMI condition as an alternative locomotion condition with an altered gain. By running sessions that deliberately interfered with the gain between wheel and VR in the probe condition (Supplementary Fig. 7, see Supplementary Text), we established that such a manipulation neither introduces remapping consistent with the removal of the locomotion signal, nor does it introduce a superposition effect. This suggests that, in this instance, the hippocampus is able to quickly correct for changes in gain and maintain a consistent representation, in agreement with existing gain modulation experiments^9,30^.

### Hippocampal CA1 representations are enriched in BMI by available signals

The evidence from these control experiments suggests an effect specific to experience of the BMI condition. Given previous findings showing that place cells fail to anchor their firing when prominent visual cues become unreliable^31–33^, we reasoned that hippocampal representations reweighted their inputs to ignore locomotion signals when they became unreliable predictors of motion in both the BMI and locomotion conditions. Extending this logic to the partial retention of representation following the BMI condition but not following the autopilot conditions, we speculated that CA1 representations incorporated a signal available during BMI conditions but not during autopilot conditions.

To test this, we used a session structure that fully excluded informative locomotion signals by using a random autopilot condition in the baseline blocks (Fig. 4a). We trained a first-generation BMI decoder on the initial autopilot block data and, as before, subsequently retrained the decoder on the first BMI block to produce a second-generation decoder for use in the second BMI block. In this configuration, we found that decoder performance was immediately accurate and BMI task performance did not meaningfully benefit from decoder retraining (Fig. 4b-c, Supplementary Fig. 8). This is consistent with our hypothesis: because the transition from the autopilot does not remove any useful signals, the established place fields (which we reasoned are exclusively anchored to visual cues) are not disrupted and widespread remapping is therefore avoided. However, while the retention of these specific place fields enabled accurate BMI decoding in the subsequent block, the population-wide CA1 responses during the BMI blocks differed strikingly. BMI-based navigation saw the emergence of additional place cells (Fig. 4d-e, an average increase of 53.6% from B1 to B2), resulting in an enriched representation consistent with the appearance of an additional source of information to integrate into the hippocampal code. This ‘autopilot + BMI’ session structure also allowed us to rule out the possibility that the superposition effect described earlier was induced purely by reward conditioning^34–36^. The new responses that formed in the BMI condition appeared primarily in the null space of the decoder weights (Fig. 4f-g). Notably, no aspects of the enriched BMI representation persisted when the animal returned to the final baseline condition (Fig. 4h-i), rendering the superposition effect exclusive to sessions combining locomotion and BMI-based navigation. This points to a potential mechanism, common to these two conditions, that drives CA1 representations.

**Fig. 4.**
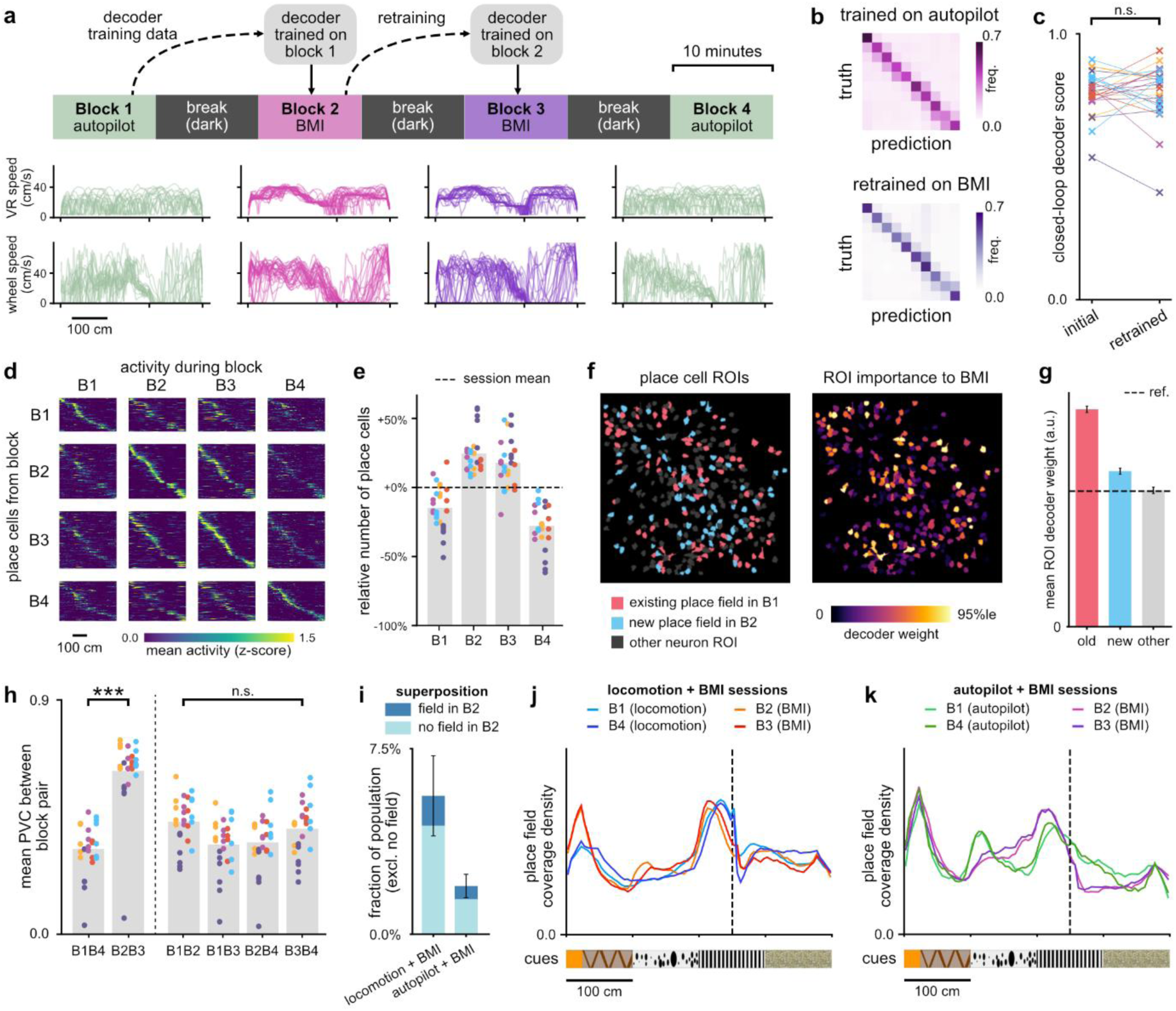
Moving from a random passive motion condition to BMI control does not induce population-wide remapping, but instead recruits additional place cells. **(a)** A modified session structure in which the baseline condition is replaced by a random-speed autopilot. **(b)** Confusion matrices aggregated over 26 all autopilot + BMI sessions evaluating closed-loop decoder performance, **(c)** alongside a comparison between overall score before and after decoder retraining for each session. **(d)** An example population ratemap from a single autopilot + BMI session, adjusted to show exclusively neurons with place fields. **(e)** The number of neurons with a place field, shown for each block in autopilot + BMI sessions, relative to the mean number of such neurons across the session. **(f)** An example imaging FoV highlighting the place fields during block 2 (BMI) of a session highlighting those neurons that retained their place field from block 1 and those that newly gained a place field, juxtaposed with an inferred decoder weight for each ROI. **(g)** These weights are higher for place cells from block 1, as their fields are visible in the training data. Bars show mean normalized decoder weight over all sessions for neurons in each category, error bars are +/- SEM. **(h)** Mean PVC between block pairs, as in Fig. 3a, for the autopilot + BMI sessions. **(i)** As in Fig. 3k, comparing the number of neurons exhibiting superposition between locomotion + BMI and autopilot + BMI sessions. **(j)** Relative coverage of the track by place fields in each block during locomotion + BMI sessions (14652 neurons). **(K)** As in (j) for the autopilot + BMI sessions (8385 neurons), illustrating a more pronounced response to the track’s visual cues.

To assess the relative impact of visual stimuli across our observed hippocampal codes, we considered the relative frequency of locations where place fields formed in locomotion, BMI, and autopilot conditions (Fig. 4j-k, Supplementary Fig. 7d). Inspecting locations near changes in wall cues, such as the 100 cm mark, revealed that autopilot-based representations anchored strongly to such stimuli, while locomotion-based ones did not. This accumulation of place cells around salient landmarks outside of the locomotion condition, as also described in earlier work^19,37^, is consistent with an active reweighting of signals available to CA1. The BMI condition exhibited a greater degree of anchoring to visual cues than the locomotion condition, but not as strongly as the autopilot condition. This suggests that the cognitive map depends only partially on visual cues during the BMI condition, and this observation is once again consistent with the presence of a signal driving CA1 that is informative exclusively in contexts in which CA1 representations are causally involved in navigation.

## Discussion

By manipulating how animals experienced navigation of a virtual environment, either by running on a wheel, from decoded activity in CA1, or as a passive observer exposed to prerecorded trajectories, we observed that representations in CA1 actively reweighted responses according to the degree and nature of their influence on navigation. Previous work has already shown that representations in CA1 are sensitive to a range of external and internal changes, including the degree of engagement with a task^17,18^, reward conditioning^37^, changes in gain between running and visual travel^9^, and time^38^.

By analyzing differences in CA1 maps during the BMI condition and other conditions, we identified partial remapping that appeared specific to the BMI manipulation and not accounted for by factors in other work. By construction, running speed is no longer navigationally informative in the BMI condition, but hippocampal activity remains relevant. BMI-specific remapping is therefore consistent with the hypothesis that, during BMI usage, CA1 reweights to depend more heavily on an internally generated signal that is present during navigation via locomotion, but absent during passive experiences of motion.

This hypothesis also provides a satisfying account for the poor first-generation BMI decoding performance and the improvement in decoding performance obtained using autopilot trial data, because the latter presumably instantiated a hippocampal representation of the environment that disregards physical movement. Such sensitivity to the context in which BMI training data were obtained contrasts strongly with BMI decoding in other brain regions, such as in the posterior parietal cortex^39^ and primary motor cortices^40,41^, that find that high offline accuracy translates to good closed-loop task performance.

Previous work using hippocampal BMIs either explicitly used rewards to condition neuron firing^35^ or transported animals directly to decoded locations^26^ by thresholding template activity patterns. Both paradigms therefore potentially mask naturally arising discrepancies between BMI and locomotion-based representations of the same environment: the former explicitly conditions BMI responses, the latter filters out responses that are inconsistent with non-BMI conditions. Indeed, there is a growing body of evidence across brain regions for response subspaces that are BMI-specific^42–44^. Notably, using differently-weighted readouts for performing the same BMI task can induce ‘memory traces’^45^, which partially resemble our observation of superposed neurons. We propose that our observed retention of place fields when moving from BMI control to locomotion, as well as the appearance of new place fields when moving to BMI control from passive motion, suggest the existence of an input signal that can actively drive CA1. This signal appears independent of self-motion or external stimuli, and it is either absent or muted when animals are unable to influence their navigation and are merely passive observers of trajectories.

While recent findings in humans suggest that hippocampal spatial representations may be volitionally evoked by ‘mental imagery’ of environments^46–49^, and while recent work in rats has described the active generation of patterns of hippocampal activity to make progress towards rewards^26^, volition remains fundamentally problematic to define in animal models. Our BMI protocol was designed not for an animal to learn how to control a new means of navigation, but instead to causally link stable hippocampal representations to stable task outcomes. The defining feature common to the BMI and to the locomotion conditions was the ability of CA1 to influence a sequence of experienced events, and therefore an explanation that does not rely on ideas of volition is the prediction of future states. When consistent neural activity within the brain generates consistent experiences in the environment, irrespective of whether this required motor activity, then the ability to embed this relationship in a cognitive map allows animals to plan. A component of a hippocampal cognitive map that distinguishes when an animal’s own actions drive outcomes, as opposed to exogenous phenomena, therefore provides a key building block for higher cognition^50^.

## Notes

## Acknowledgements

We are grateful to Marius Bauza for his development of the VR environment; Stella Ma, Pauline Kerekes, and Marino Krstulovic for making available imaging datasets instrumental to the design and testing of the BMI; Kou Huang for a comparative evaluation of place cell identification algorithms; and Ethan Sorrell and Dan Wilson for sharing their expertise in optical BMI experiments.

## Data availability

All data needed to assess the findings, code for analysis described in the Methods, and code for our custom BMI software are available on Zenodo^51^.

## Methods

### Animal handling and surgical procedures

All experimental procedures were conducted in accordance with the UK Animals (Scientific Procedures) Act 1986 (ASPA), following ethical review by the University of Cambridge Animal Welfare and Ethical Review Body (AWERB). All procedures were performed by trained personnel authorized under Home Office Personal and Project licenses.

C57BL/6J wildtype male mice were obtained from Charles River and a total of 11 mice were used for experiments. Mice were housed in conventional plastic cages (16 cm x 27 cm x 18 cm) and were group-housed where possible. Water was supplied ad libitum. Modest food restriction was used during experimental periods to encourage running and reward-seeking behavior. All mice were kept on a 12:12 hour light:dark cycle in an animal facility with regulated temperature (21-23°C) and humidity (50-60%).

Viral injection surgery took place between 58 and 65 days of age. The implantation of an imaging cannula for optical access was done in two steps: 1) surgery for viral injection in CA1 region and head-plate implantation; 2) surgery for implanting a small imaging cannula for 2-photon imaging. Mice were anaesthetized with 1-3% isoflurane in O_2_ and placed in a stereotaxic frame. The scalp was shaved and swabbed with iodine before incisions were made to expose bregma, from which coordinates were measured. Analgesic (Metacam, 0.05 mg/10g body weight) and antibiotics (Baytril, 0.05/10g) were administered via subcutaneous injection. Body temperature was maintained at around 35°C and breathing was monitored throughout surgery. Ointment was applied to protect the eyes from dehydration and scarring. Mice were placed in a heated chamber for 1-2 hours following surgery to aid recovery before transfer back to their home-cage. Post-operative care, which generally lasted 5 days, provided Metacam and Baytril orally and monitored body weight.

The head plate consisted of a custom-made Aluminum piece with a central opening for imaging, flanked by two ‘wings’ which could be fastened with screws to fixed points relative to the microscope objective during experiments. Outside of experiments, the central opening was kept covered by a screw cap to keep the imaging cannula clean. Viral injection was made in the right hemisphere, and two contralateral sites of craniotomy were drilled for insertion of jeweler’s screws to provide additional support for the head-plate. Opaque dental cement (Superbond C&B) was applied to permanently attach the head-plate to the skull. A Nanofil needle and syringe (World Precision Instruments) controlled by a micropump were employed for viral injection. A craniotomy of approximately 1 mm was made atop the injection site with the following coordinates: AP: 2.0 mm posterior from Bregma; ML: 1.8 mm from midline; DV: 1.3 mm beneath dura^52^. To express Calcium indicator in the dorsal hippocampus stratum pyramidale, 200 nL of titre 2.3 x 10^13^ GC/mL AAV1-syn-jGCaMP8f-WPRE-SV40 (Addgene 162376-AAV1)^53^ diluted 1:1 in artificial cerebrospinal fluid was injected at 30 nL/min, followed by a 10-minute wait to minimize backflow before retracting the needle. Exposed skull was covered with Kwik-cast and the head-plate covered with a screw cap.

One week after viral injection, a second surgery implanted the imaging cannula (outer diameter: 2.7 mm) with a thin circular glass window attached to its bottom over the injection site, as previously described by Dombeck and colleagues^52^.

### Animal training in the VR environment

We used a custom-built closed-loop 1D virtual reality (VR) environment, with rendering implemented in Unity and synchronization with both a rotary encoder on the wheel (to track running speed) and the microscope mirror galvanometer (to synchronize imaging frames to VR state) implemented in LabView. The VR environment was displayed across two monitors mounted facing the Styrofoam wheel to provide a wide field of view.

Following post-operative recovery, mice were handled and habituated on the Styrofoam wheel in the VR setup for two days. Mice were then placed on food restriction and their health and body weight were checked daily. The body weight was kept at no lighter than 85% of the original weight. Mice were then subjected to head-fixing, initially for several minutes, with duration increasing in subsequent sessions until they were able to be head-fixed for 30-40 minutes. While head-fixed, mice were encouraged to run on the wheel and received rewards for doing so. During this habituation, the monitors displayed a dark screen. The reward location on the track as randomly distributed to prevent the mice from developing anticipatory licking behavior based on distance, as well as to encourage consistent exploratory licking of the reward spout. Once a mouse could complete 100 laps of the 400 cm track, the animal was introduced to the VR environment (screens no longer dark, reward at fixed location) on the following session. This habituation and training program took an average of 1-2 weeks to complete.

When first introduced to the cue-rich VR track, mice displayed scattered licking along the track, as they were unfamiliar with the environment and reward location. Following several days of exposure, mice began to exhibit behavior associated with reward anticipation, exemplified by improved licking specificity and deceleration as the animal approached the reward site. A mouse was deemed well-trained when anticipatory licking behavior became stable and the running velocity profiles were comparable on two consecutive days. Typically, it took mice 5-7 days to become experts.

### 2-photon imaging hardware

The imaging microscope was supplied and built by Bruker Corporation, including a Ti:sapphire excitation laser (Chameleon Ultra II, Coherent) with a frequency of 920 nm and 60-70 mW average power at FoV, and mounted on an antivibration table (Thorlabs Inc.). A LWD water dipping 16x 0.8 NA objective (Nikon) was used, and the cannula was filled with distilled water for imaging. We used Prairie View 5.6 (Bruker Corporation) for software control of the microscope, either manually through its Graphical User Interface, or programmatically via an interface with our in-house BMI software, ReTiCaDe^51^. We blocked light leakage from the VR screens to the objective lens with a removable custom shield of black polyurethane-coated nylon fabric fitted to the head-plate. Imaging sites were coarsely located using epifluorescence prior to more exact alignment using 2-photon imaging.

### Animal screening for progression to BMI sessions

After animals became experts on the task, which typically coincided with good indicator expression, we screened animals for progression to BMI based on imaging quality. We imaged 10-minute blocks, similar to those performed at the start of a BMI experimental session to provide training data for decoders, and inspected the imaging to verify:

- Minimal tissue movement in the z-axis during running, as this is not feasible to compensate for with planar motion correction algorithms
- Good indicator expression within dorsal CA1 hippocampus stratum pyramidale, with an excess of 150 neuron ROIs in the FoV extracted with suite2p^54^
- Sufficient presence of place cells in the population, with at least 15% of imaged cells forming a place field

We evaluated offline open-loop BMI decoder performance on a withheld number of laps from this session before proceeding to closed-loop BMI sessions, but this was not used as a selection criterion, as the criteria above were sufficient to ensure accurate open-loop BMI in all cases.

### Experimental schedule

The experimental schedule, which describes which animals performed which types of session on any given day, is shown in Supplementary Fig. 3b. Of the 11 animals, the first animal (animal 01) was also used for a series of technical systems checks, and so experienced a variety of non-standard configurations of the VR and BMI environment. It was therefore excluded from analyses relating to changes in BMI decoder performance between sessions. The remaining 10 animals were divided into two cohorts of 5, with the first cohort (animals 02 through 06) experiencing several consecutive locomotion + BMI sessions and the second cohort experiencing several consecutive autopilot + BMI sessions.

### Control of VR navigation from real-time decoding of 2-photon imaging

The imaging FoV was scanned in a serpentine pattern, spanning the entirety of the FoV at 30 Hz and acquiring a 512 x 512 pixel image with 3 samples per pixel. The raw stream of data arriving from the microscope was marshalled into a ring-buffer in shared memory, from which the latest full frame was asynchronously read-out by our custom signal processing pipeline and neural decoder. The output of the decoder, corresponding to a control signal to set forward velocity in the simulation of a VR environment, was broadcast to a secondary computer responsible for rendering and orchestrating the VR task. Measurements of the galvanometer controlling the laser mirror for the microscope were taken directly from the VR task computer via a side-channel that bypassed the imaging computer to ensure that individual image frames could be exactly synchronized with the visually-rendered VR scene during analysis (Supplementary Fig. 1a).

### Construction of decoder feature space via 2-photon signal-processing

Instead of producing individual ROI masks to act as the input to the decoder, as has been previously demonstrated in offline validation^55,56^ of optical BMI in CA1, we developed a signal processing pipeline that transformed raw images into a 32x32 feature space in which each pixel was a close surrogate to underlying neuron ROIs (Supplementary Fig. 1b). This constant input vector size led to consistent decoder training times and, combined with not needing to extract individual ROIs, short and consistent processing times between ingestion of image data and resulting motion in the VR environment. It consisted of the following steps:

- Low-pass filtering the 512x512 pixel image with a 2D Gaussian smoothing filter and downsampling the image resolution to 128x128
- Applying non-rigid motion correction relative to a reference frame
- Applying a real-time dF/F filter to each pixel
- Applying a spatial band-pass filter to the image
- Thresholding the image to include positive values only
- Downsampling to a 32x32 pixel image to produce the feature space

The 2D Gaussian smoothing filter used a standard deviation parameter sigma σ equivalent to 9.6μm (accounting for the image resolution at the point the filter is applied).

The non-rigid motion correction used the scikit-image implementation^57^ of the Iterative Lucas-Kanade optical flow correction algorithm^58^, with the maximum window evaluated around each pixel using a radius equivalent to 28μm, and the evaluation of a single warping, as displacements were typically small and each subsequent warping required trading off against computational cost. Each motion-corrected frame was aligned against a reference frame. For closed-loop BMI decoding, we constructed a reference frame from the mean pixel-wise fluorescence over the 10-minute block used as training data for the decoder.

The real-time dF/F filter adapts the typical formulation of a dF/F filter (which normalizes by mean fluorescence over an entire block duration) to instead use an exponentially smoothed moving average, as has been previously used in optical BMI designs^39^. This produces a filtered signal *F*^′^(*t*) from a raw fluorescence signal *F*(*t*).

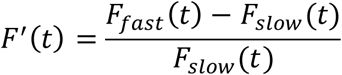

Where:

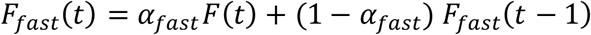

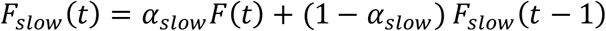

The constants *α*_*fast*_ and *α*_*slow*_ were set to 0.3 and 10^-3^ respectively, which correspond to time constants of 33.3 s and 0.09 s respectively. Due to the relatively long time constant for *F*_*slow*_(*t*), we ‘burned in’ the filter on a stream of frames to produce an unbiased initial state. In offline analysis, the filter was burned in by first ingesting *F*(*t*) in reverse time order. For online, closed-loop BMI blocks, we initialized both *F*_*fast*_(*t*) and *F*_*slow*_(*t*) to the mean fluorescence value for the relevant pixel over the course of the training data.

The spatial band-pass filter applied after temporal filtering used a 2D difference-of-Gaussians parameterized by σ_lower_=20μm and σ_upper_=4μm.

### Real-time classification of track position

The track was divided into 10 spatial bins. This number of bins was chosen such that the bins did not always align with changes in wall cues, to ensure that the reward site was well-contained within a single bin, and because it was feasible to decode accurately to this resolution with 10 minutes of training data. To account for uneven occupancy of spatial bins, training data classes were balanced prior to model fitting by weighting samples to ensure equal total sample weight in each bin.

The SVM classifier used an L1 penalty to provide sparse regularization, discouraging allocation of non-zero weights to features that provided minimal information about track location. The hyperparameter controlling the strength of the regularization (denoted *C* in most texts) was set to 0.2. This was initially selected by a sweep of the parameter-space evaluating performance on unseen laps from the offline evaluation sessions and subsequently fixed for the use of all closed-loop BMI sessions.

In addition to the SVM classifier used for the BMI described in the main text, we also compared the offline performance of two other classifiers (see Supplementary Text). The Naïve Bayes classifier assumed feature likelihood to be Gaussian, estimating a mean and variance for each neuron in each bin. The feedforward neural network was a simple multi-layer perceptron, parameterized by the size of its single hidden layer (selected through manual trial and error on offline data). We selected the best-performing network as a comparison point. The network presented in the supplement used a tanh activation function, was fully connected, used early stopping, and was trained with gradient descent (Adam). It used L2 regularization, with the strength of the regularization (denoted α in most texts) determined by 5-fold cross-validation. All classifiers used existing implementations from the scikit-learn Python package^59^.

### Mapping of decoded spatial bin to forward velocity

We transformed the output of the decoder’s classifier (the identity of a spatial bin) to forward velocity through the VR environment by selecting the corresponding forward velocity from a mapping. This mapping was empirically constructed by inspection of stereotyped performance of the VR task, such that when the decoder output corresponds to a spatial bin in which animals typically run fast the BMI-controlled velocity will be high, and when the decoder output corresponds to a spatial bin in which animals are typically slow, the BMI-controlled velocity will be slow. The highest achievable velocity under BMI control was 40 cm/s and the lowest achievable velocity was 4 cm/s (set to a small non-zero value in order to prevent the animal becoming stuck). The same mapping was used for all animals in all BMI-controlled sessions.

In order to avoid visually jarring switches between speeds in the event of noisy decoder output, we limited the maximum acceleration during BMI control to 80 cm/s^2^, which was approximately double the maximum acceleration we observed animals performing when running on the wheel.

### Decoder accuracy evaluation

We produced confusion matrices to visualize the accuracy of decoders. These compare the output of the decoder, a predicted spatial bin, to the ground truth spatial bin at each imaging frame. Each row of the matrix represents occurrences of the ground truth (instances of the animal being within the bin) and each column represents occurrences of particular decoder outputs. Values in individual cells represent relative rates of occurrence, normalized by the ground truth (such that each row in the matrix sums to unity).

These confusion matrices were simplified into a single value score from 0.0 (no accuracy) to 1.0 (perfect accuracy). First, we defined the decoder score for each spatial bin as the fraction of samples that were classified either completely correctly or to within one neighboring bin, similarly to Chen et al.^55^. The overall decoder score was then computed as the median of the bin scores. This median-over-bins scoring strategy addresses sampling bias issues that arise in closed-loop contexts in which decoder output can directly influence which spatial bins are over- or under-sampled.

### Comparison between feature space and neuron-based decoding

In offline analysis comparing decoder performance between relying on the feature space and on neuron ROIs, we used the same SVM classifier as described above, with the following changes:

- The input to the SVM was the dF/F filtered traces of all neuron ROIs
- To account for differences in the input dimension, we re-swept values for the hyperparameter C to find a value appropriate for the ROI inputs (eventually using C=0.05)

### Extraction of decoder neuron weights from feature weights

To infer the weight allocated to a given neuron for a set of feature weights, we iterated over each pixel in the neuron’s footprint (at a resolution of 512x512 pixels) to determine the proportion of overlap with each feature in the feature space (32x32 pixels). We calculated a ‘raw’ weight for each bin for each neuron ROI by performing an average of the weights in the feature space, with the averaging weighted by the proportion of overlap by the ROI. We then normalized these raw weights across all neuron ROIs within each class (spatial bin) by dividing by the standard deviation of raw weights within that class.

The inferred per-neuron decoder weights can therefore be displayed by spatial bin (as in Fig. 1d). When we present them as aggregated (as in Fig. 4f-g), we take each neuron’s largest weight across bins. The thresholding step of the signal-processing pipeline results in weights being (nearly entirely) positive.

### Passive motion conditions

We produced three different passive motion (‘autopilot’) conditions as controls. The ‘constant’ autopilot condition fixed the animal’s VR speed at 30 cm/s (Supplementary Fig. 6c). The ‘naturalistic’ condition replaced the output of the decoder with the animal’s immediate ground-truth position on the track before mapping decoded spatial bin to forward speed, and thereby passively produced well-executed laps (Supplementary Fig. 6e).

The ‘random’ condition was experienced by animals 07-11 and was generated by taking existing BMI traversals performed by animals 02-06 from block 2 (i.e. poor decoder performance). These existing traversals were then temporally shifted by a random offset between 10% and 90% of the block duration (uniform sampling) and then played back to an animal during the random autopilot condition. This ensured that the distribution of forward velocities and the number of laps experienced by the animals during the random autopilot condition matched the distributions of velocities and laps from typical block 2 laps during locomotion + BMI sessions. No animal was shown the same ‘replay’ more than once, preventing animals from learning any temporal structure of the passive motion.

### Gain modulation condition

The gain modulation condition used as an additional control (Supplementary Fig. 7e) altered the ratio between rotation of the wheel and distance travelled on the VR track. Instead of a constant gain alteration, we implemented a system for altering the gain as a function of track location. We used the same 40 cm spatial bins used for decoder classification and implemented a gain profile that attenuated speed in the typical region of fast running and amplified speed in the typical region of slow running (exact gain profile shown in Supplementary Fig. 7f).

### Offline extraction of neuron ROI fluorescence

ROIs were extracted from imaging sessions using suite2p^54^, treating only those ROIs marked as cells with high confidence by suite2p as ‘neuron ROIs’. Neuropil fluorescence at each timestep was subtracted from the fluorescence traces for each neuron ROI to produce a corrected fluorescence:

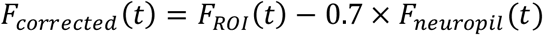

These corrected traces were then dF/F filtered using the same real-time dF/F filter described above. In instances where we use or display z-scores of traces, we evaluated the mean and standard deviation of the dF/F traces of each neuron over all four blocks of a session to produce a per-neuron z-score trace.

### Construction of ratemaps

We constructed ratemaps at a resolution of 5 cm spatial bins on the track. To compute the ratemap for an individual neuron ROI within a single block, we evaluated the mean z-scored dF/F value within each of those bins (the sum of z-scored dF/F values within each bin divided by the animal’s dwell time within that bin).

When displaying ratemaps across entire populations and 4 blocks, we first construct a neuron ordering per block based on which spatial bin (from start to end of the track) contains that neuron’s largest ratemap value within that block. To produce the 4x4 grid, we re-plot all neuron ratemaps in each block according to each block ordering.

### Place cell identification

Place cells were identified from the processed dF/F signals for each neuron ROI according to the following steps, which were adapted to our task and dataset from the Combination Method described by Grijseels et al.^60^. The first step of this method requires identifying period of ‘Calcium events’, which are spikes in Calcium activity defined as:

- Starting at the point in time at which activity exceeds the upper threshold T_upper_
- Ending at the point in time at which activity falls below the lower threshold T_lower_
- Lasting for at least 3 frames (∼100 ms)

The two thresholds are defined as:

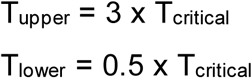

Where T_critical_ is chosen for a neuron ROI by slicing a block timeseries into 2-second windows, evaluating the standard deviation of the dF/F activity in each window, and then taking the median of these standard deviations over all 2-second windows.

The second step determines candidate regions for place fields. We computed a ratemap for each ROI at a resolution of 5 cm. We then defined a cut-off threshold Y such that Y=0.25(p_max_-p_25_), where p_25_ is the 25^th^ percentile of the ratemap bins and p_max_ is the largest ratemap bin value. Candidate place fields are contiguous regions of the ratemap exceeding the cut-off threshold Y, at least 15 cm in length, and no more than 200 cm in length. We applied two initial criteria to determine whether to retain candidate place fields:

- The mean ratemap value within the candidate field must exceed 4x the mean ratemap outside value outside of all candidate fields for the ROI
- The ‘activity ratio’ must exceed 25%. The activity ratio is the proportion of time classified as a Calcium event while the animal is within the spatial bounds of the place field

All ROIs not meeting these criteria were discarded as candidate place fields.

Finally, we performed a comparison against shuffles of neural activity relative to track position. This broke the neuron ROI’s dF/F timeseries into blocks of 120 frames (∼5 seconds) and shuffled the order of the blocks while keeping the timeseries of the animal’s position unaltered. We then performed the same place-field identification procedure as above and rejected any candidate fields that that did not exceed an activity ratio within the field greater than 95% of 1,000 evaluations on different random shuffles of the data.

While it is common in other work performing place field identification to remove samples in which the animal is stationary (e.g. moving at < 5 cm/s), we chose not to remove these samples as our experimental setup effectively decouples running from traversal of the VR environment.

### Relative change in place cell counts

For *n*_*i*_ place cells in the i^th^ block within a single session, we define the proportional change in place cell count relative to the session mean as:

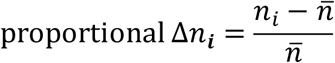

Where *n̅* is the mean number of place cells over all blocks in that session.

### Population vector correlations

PVC is a measure of similarity of population activity between a pair of blocks at a given location. We first computed the neuron ratemaps for each block, as described above. This produces an N x M matrix for each block (N neurons, M spatial bins in the ratemap). The PVC between two blocks at a given location, corresponding to the j ^th^ spatial bin, is the Pearson correlation coefficient between the two j^th^ columns of the matrices for each block. We do not make use of PVC at any specific location in this work, but instead report the mean PVC over all bins as an overall measure of similarity.

### PVC evaluations on sub-ratemaps

For Supplementary Fig. 5, we used mean PVC to evaluate the effects of correlation between locomotion speed and VR movement during BMI (Supplementary Fig. 5b), time (Supplementary Fig. 5c), and the presence of cells with ‘superposed’ fields (Supplementary Fig. 5d).

In the first case, we were interested in representational similarity *within* a single block. For block 3 in each session, we isolated each lap of the track and then ranked all the laps according to the Pearson correlation between wheel speed and VR speed. We then constructed two sub-ratemaps, one for the upper 50% of ranked laps, and one for the lower 50% of ranked laps, and evaluated the mean PVC between these two sub-ratemaps. To provide comparison points, we repeated the process of lap splitting in two additional ways: by taking alternating laps (resulting in a group of odd laps and a group of even laps) and by taking the first half and second half of laps separately. In the second case, we were interested in whether the representational similarity between Block 3 and Block 4 during BMI was merely transient and would disappear after the animal experienced several laps of locomotion control during Block 4. Here, we took Block 4 and divided its laps to construct sub-ratemaps according to four different schemes: odd laps, even laps, the first 50% of laps, and the last 50% of laps. We then evaluated the mean PVC between the complete ratemap for Block 3 and each sub-ratemap for block 4. The comparison of the first 50% to the last 50% serves as a measure of transience, while the odd to even comparison provides a baseline. We showed this comparison for locomotion + BMI sessions, but also for locomotion + locomotion sessions in order to illustrate the effect of natural representational turnover over the course of block 4 unrelated to superposed representations.

In the third case, we were interested in the extent to which the cells that we heuristically categorized as superposed were responsible for the discrepancy between the mean PVC between B1B2 and B3B4 during locomotion + BMI sessions. Here, we evaluated the mean PVC between these block pairings after excluding cells meeting these criteria from the ratemaps.

### Place field coverage density

To estimate place field coverage of the track, we first discretize the track at the same resolution as place cell identification and ratemap construction (5cm) and then mark each of these bins has having initially zero coverage. For each neuron, we increment each bin’s coverage value at each bin overlapped by that neuron’s place fields. After all neurons have contributed to the coverage map, we normalize the coverage map to have unit area so that it represents coverage density.

### Heuristic categorization of neuron responses

We applied the following criteria to automatically categorize neurons based on differences between place field presence and location across a 4-block session with an initial reference block, two probe blocks, and a final reference block. These criteria were applied sequentially, halting once a neuron was successfully categorized.

1. Complete absence of place fields in all blocks: not a place cell
2. Exactly one block contains place fields: single-block transient field
3. All blocks have at least one place field that matches at least one place field in every other block: stable
4. Blocks 1 and 4 share a field and no other blocks share a field: modulated, reference-specific
5. Blocks 2 and 3 share a field and no other blocks share a field: modulated, probe-specific
6. Blocks 1 and 4 share a field and blocks 2 and 3 share a field, but no fields are shared between any other pair of blocks: modulated, field moves during probe
7. Blocks 1 and 4 share a field and blocks 2, 3, and 4 share a field: superposition, 1&4+3&4
8. Blocks 1 and 4 share a field and blocks 3 and 4 share a field: superposition, 1&4+3&4
9. Other configurations in which exactly two blocks share a field: other, pair
10. Other configurations in which a subgroup of three blocks all share a field and the remaining block shares no fields with any blocks: other, triad
11. Unclassified

Place fields identified across two blocks must satisfy either of the following criteria in order to qualify as ‘matched’ or ‘shared’:

- The length of the track overlapped by the two fields is at least half the length of the smaller of the two fields
- The field centers are no more than 15 cm apart

For Supplementary Fig. 5d, we additionally considered a ‘relaxed’ definition of superposed representations that did not explicitly require a distinct field in Block 1 (i.e. a superposition with the empty map). For this relaxed definition, we included cells from the ‘Other’ category corresponding to a configuration in which exactly Blocks 2, 3, and 4 or exactly Blocks 3 and 4 shared a common field.

### VR and wheel speeds

When wheel speed or VR speed are plotted showing multiple laps on the same axes, as in Fig. 2a, Fig. 4a, Supplementary Fig. 5, and Supplementary Fig. 7, the traces are smoothed with a boxcar filter (zero-padded between laps, 500 ms window) for ease of viewing. This smoothing is not applied when the laps are viewed individually, as in Fig. 2j-k.

We compute both the Pearson correlation coefficient between the wheel speed and the target trajectory (as shown in Fig. 2l) and between the wheel speed and the animal’s actual motion through the VR environment (as shown in Supplementary Fig. 4e), restricting this calculation to the periods of time in which the animal is in the VR environment (i.e. excluding the inter-lap darkness interval).

### Anticipatory licking scoring

To score the accuracy of anticipatory licking behavior, we first categorize each lick event based on its track position into one of three categories:

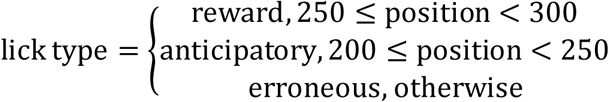

For each lap of track, we calculate the fraction of anticipatory licking events among both anticipatory and erroneous licking events.

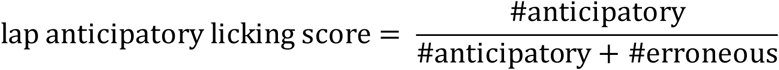

The anticipatory licking score for an entire block is the mean of this score over all laps.

### Engagement metric

We combine the anticipatory licking score and the correlation between wheel speed and VR speed into a single metric. We perform this by computing the z-score for each metric over all considered blocks and then defining the engagement metric as the larger of the two z-scores for that trial. The rationale for this combination is that the task can be well-executed even if one of these metrics in isolation is low, but that animals scoring poorly on both metrics are more likely to be disengaged.

### Statistical methods for PVC similarity comparisons

We treat the six different block pairings for which we evaluate mean PVC by first splitting them into two categories: pairings in which the control modalities match (baseline vs baseline and probe vs probe) and pairings in which they do not (baseline vs probe). We then apply three statistical tests.

The first test checks whether PVCs evaluated in same-modality pairings are different from those evaluated in mixed modality pairings. This is accomplished with a two-sided t-test that treats animal identity as a subject identifier and the mean PVC as the within-subjects factor.

The second test checks whether the two trial pairings in the same-modality group (baseline vs baseline and probe vs probe) have significantly different PVCs. This is accomplished with a paired (by session) two-sided t-test that treats animal identity as a subject identifier and the mean PVC as the within-subjects factor.

The third test checks whether experience of the probe condition alters the baseline representation in a way that makes it more similar to the probe condition. This is accomplished by comparing the B1B2 pairing to the B3B4 pairing, and takes place in two stages. First, we perform a repeated-measures ANOVA among all block pairings of differing modality (B1B2, B1B3, B2B4, and B3B4), treating animal identity as a subject identifier and the mean PVC as the within-subjects faster. After confirming the ANOVA result, we then perform pairwise comparisons between block pairings with paired (by session) two-sided t-tests (also treating animal identity as a subject identifier and mean PVC as the within-subjects factor) and use a Bonferroni correction to account for the multiple comparisons. We then report the p-value associated with the B1B2 vs B3B4 test.

These tests are all implemented using the reference implementation from the Pingouin statistics library^61^, and full results are tabulated in Supplementary Table 1.

### Statistical methods for linear regressions

For the linear regressions used in Fig. 2i, Fig. 2l, and Supplementary Fig. 4c-d, we fit an ordinary least-squares model to the data. To compute the p-values corrected for animal identity, we used a linear mixed-effects model using a reference implementation from the statsmodels Python library^62^, which treats each animal as having its own y-intercept in the linear model.

For the linear regressions in Supplementary Fig. 4e-f, we followed the same procedure, but we also showed an additional fit with outliers removed. To remove outliers in these plots, we first computed the ordinary least squares regression. We then removed all samples with squared residuals above the 95^th^ percentile and refit to the remaining samples.

### Statistical methods for comparison of changes in decoder score post-retraining

When comparing decoder score during block 2 (pre-retraining) and block 3 (post-retraining), as in Fig. 2g and Fig. 4c, we used a paired (same session) single-sided t-test, using a reference implementation from the SciPy library^63^.

For Supplementary Fig. 8c, we also aimed to compare BMI scores between block 2 and block 3, but we are also dealing with two different cohorts of 5 animals each. In the case of comparing between cohorts, comparisons cannot be paired within session, and so for consistency we used two-sided independent t-tests in all comparisons made for this figure.

### Statistical methods for differences in anticipatory licking behavior

We used a repeated measures ANOVA, treating the licking score as the dependent variable, block index as the within-subjects factor, and animal identity as the subject identifier, using a reference implementation from the Pingouin software library^61^.

## Supplementary material

### Supplementary text

#### Additional control sessions

In addition to the three session types described in the main text (locomotion + BMI, autopilot + BMI, and three varieties of locomotion + autopilot), we ran two other major session types as controls: locomotion + locomotion and locomotion + gain modulation.

In locomotion + locomotion sessions, all 4 blocks consisted of the locomotion condition (Supplementary Fig. 7a). This allowed us to sanity-check that no dramatic remapping of the same environment took place due to the 10-minute darkness period between blocks and allowed us to quantify the extent of baseline representational turnover from one block to the next (Supplementary Fig. 7b). As previously reported^38,52^, gradual representational turnover meant that blocks further apart in time were less similar to each other: this is why we chose to compare the temporally adjacent pairings B1B2 and B3B4 when quantifying the superposition effect with mean PVC. We also use these control sessions to highlight that the density of place cells in the population gradually decreases as the session progresses, likely due to reducing novelty or engagement, and so do not find this effect noteworthy in our locomotion + BMI sessions (Fig. 3b). Despite the small decrease in place cell numbers, the locations that remain salient on the track remain unchanged, with the distribution of place cells remaining consistent from block to block (Supplementary Fig. 7d).

In locomotion + gain modulation sessions, we used a probe condition in which the animal’s locomotion was still causally responsible for its travel along the track, but we perturbed the ‘gain’ of its running by altering the amount of travel through the VR environment produced by a single rotation of the wheel (Supplementary Fig. 7e). This gain modulation was applied at varying intensities as a function of location on the track, either attenuating or exaggerating the motion by a factor of up to 2 (Supplementary Fig. 7f). We used this control session to account for a potential confound: we used the same target BMI trajectory for all animals, despite some animal-to-animal variation in their average running speeds. What if, during the BMI condition, the animal believed its running on the wheel was still responsible for its motion through VR, only with a different gain? We found that, unlike the transition from locomotion to BMI, the transition from locomotion to the gain modulated condition did not induce a population-wide remapping, nor did it introduce a superposition effect (Supplementary Fig. 7g-h).

#### Technical details of the BMI design

With our design, the mean time between a neuron in the population spiking and a visible change on the VR screen was slightly less than 60 ms (Supplementary Fig. 1a). Approximately 30 ms of this delay was due to refresh rate of the monitor and the scanning of the 2-photon image. A further ∼20 ms was taken up by buffering data in transit. Signal processing and decoding themselves only took ∼9 ms, and over 80% of this computation time was dedicated to applying motion correction. We were able to empirically verify total latencies via a side-channel: the computer responsible for rendering the VR environment also directly read out from the galvanometer controlling the laser mirrors, allowing us to detect each new imaging frame and align it to the VR environment state during analysis.

The signal processing pipeline used a thresholding step (see Methods) in the construction of the feature space (Supplementary Fig. 1b). This meant that all activity below a threshold (effectively mean activity) was set to zero. This nonlinearity meant that, for each bin, the decoder would learn only to depend on the activity of a feature (rather than its absence), making the decoder weights interpretable by making them correspond to features that were active during the relevant spatial bin on the track, and hence to correlate closely with place cells.

The ‘failure mode’ of the decoder during pre-retraining closed-loop blocks was to overpredict the bin containing the reward site (see the vertical column of the confusion matrix in Fig. 2e). While this bin was the most frequently sampled in the training data, the training of the decoder used class-balancing to correct for unbalanced sampling (see Methods). Instead, this tendency of the decoder to predict the reward site following a change in underlying representation is a result of which parts of the track form more place cells. The reward site is a particularly salient location and attracts many more place cells (Supplementary fig. 7d, Fig. 4j-k), which results in a greater number of decoder features with non-zero weights for that bin (Supplementary Fig. 2a). We validated this hypothesis by inspecting, post-hoc, the output of the decoder when each feature was replaced by independent Gaussian noise (Supplementary Fig. 2b), and finding that this also produced overprediction of the reward site. Should future BMI work find it important to have decoder failure modes that generate uniform mispredictions, rather than mispredictions that bias towards a specific bin, this could be achieved by feature rebalancing across bins. In our case, we found this explicit failure mode convenient, both because it was an immediately obvious indicator of when the decoder was not performing well in closed-loop, and because the bin mapped to the slowest forward velocity via the location-to-speed mapping.

During design of the BMI, we considered a variety of classifier designs for the decoder. In addition to the SVM-based classifier, we also evaluated feedforward neural networks with a single hidden layer (as an example of a classifier that is capable of learning more complex encodings than a linear classifier) and a Naïve Bayes decoder (as a reference point, as this choice of decoder is common in the hippocampal literature). The Naïve Bayes decoder was considerably less accurate, but the feedforward neural network and the SVM classifiers performed nearly identically (Supplementary Fig. 2c-d). This is consistent with existing studies of hippocampal decoding from Calcium imaging data^56^, and is a result of representations in CA1 being sparse and therefore well-suited to linear decision boundaries.

Finally, we used evaluations of offline decoding performance on the same data to compare our feature space, which could easily be constructed in real-time for closed-loop BMI, to decoding from the fluorescence of neuron ROIs recovered via suite2p post-hoc. We found that their decoder accuracies were essentially identical (Supplementary Fig. 2e), strongly implying that they contained the same information. However, as the feature space is not guaranteed to exactly line up with the footprints of neuron ROIs (Supplementary Fig. 2g), it is possible for neighboring pixels in the feature space to be assigned small decoder weights (Supplementary Fig. 2h-i), and this may therefore (in post-hoc analysis) spuriously assign some amount of ‘decoder importance’ to nearby a spatially uninformative neuron ROI that abuts an informative neuron. For this reason, when we perform analysis of decoder weights, we compare the inferred weights of neuron ROIs relative to each other, rather than to zero (as in Fig. 4g).

#### Factors affecting closed-loop BMI performance

We described earlier, in Fig. 2g and Fig. 4c, the accuracy of the decoder during closed-loop control of the task via BMI. While there was an obvious improvement in accuracy when decoders trained on locomotion blocks were retrained on BMI blocks, there was still considerable variance in the retrained decoder score between sessions (Supplementary Fig. 4a-b). In the main text, we explored an improvement with experience in the BMI condition based on increasing density of place cells in the population. Not all animals had the same number of ROIs visible in their FoV (Supplementary Fig. 3a), meaning that some had richer input feature spaces to decode from. Nevertheless, it is the number of place cells in the population, rather than the size of the imaged population, that primarily governs decoder accuracy (Supplementary Fig. 4c-d).

The density of place cells that occurs in a population depends strongly on the extent of the animal’s engagement with a spatial task^17^. We considered two behavioral readouts from the animal to approximate a level of engagement with the task: wheel manipulation and anticipatory licking. In isolation, these readouts give us limited information (Supplementary Fig. 4e-f): for example, we have ample instances of animals performing well at the task despite minimal agreement between wheel manipulation and VR movement (Supplementary Fig. 4e displays a positive correlation largely because the bottom right quadrant, strong agreement between wheel manipulation and VR movement despite low decoder accuracy, is not realistically reachable). Nevertheless, if we combine these two readouts into an overall relative measure of task engagement (see Methods), we find that engagement is well-correlated with decoder accuracy during block 3 (Supplementary Fig. 4g).

While animals gained experience with the locomotion + BMI sessions improved their BMI performance over time, we found that a cohort of 5 animals performing the autopilot + BMI sessions for several consecutive days did not experience meaningful changes in the density of place cells (Supplementary Fig. S8a). Instead, we found that there were sufficiently many place cells on the first day to achieve good BMI performance (Supplementary Fig. 8b). This contrast suggests that what is improving with experience for the locomotion + BMI cohort is the ability to simultaneously maintain two separate representations for locomotion and BMI. After the autopilot + BMI cohort had performed several consecutive sessions (Supplementary Fig. 3b), we exposed them to the locomotion + BMI session type. We then compared performance between the two cohorts, restricting our analysis to sessions in which the locomotion + BMI cohort were experienced with the BMI condition (at least 10 days following the first experience with BMI) in order to account for their improvement in performance over time. We found that BMI decoder accuracy during block 3 (post-decoder retraining) was not significantly different between cohorts (Supplementary Fig. 8c). However, the cohort with experience in the autopilot + BMI condition had significantly higher decoder accuracies during block 2 (pre-decoder retraining).

## Supplementary Display Items

**Supplementary Fig. 1.**
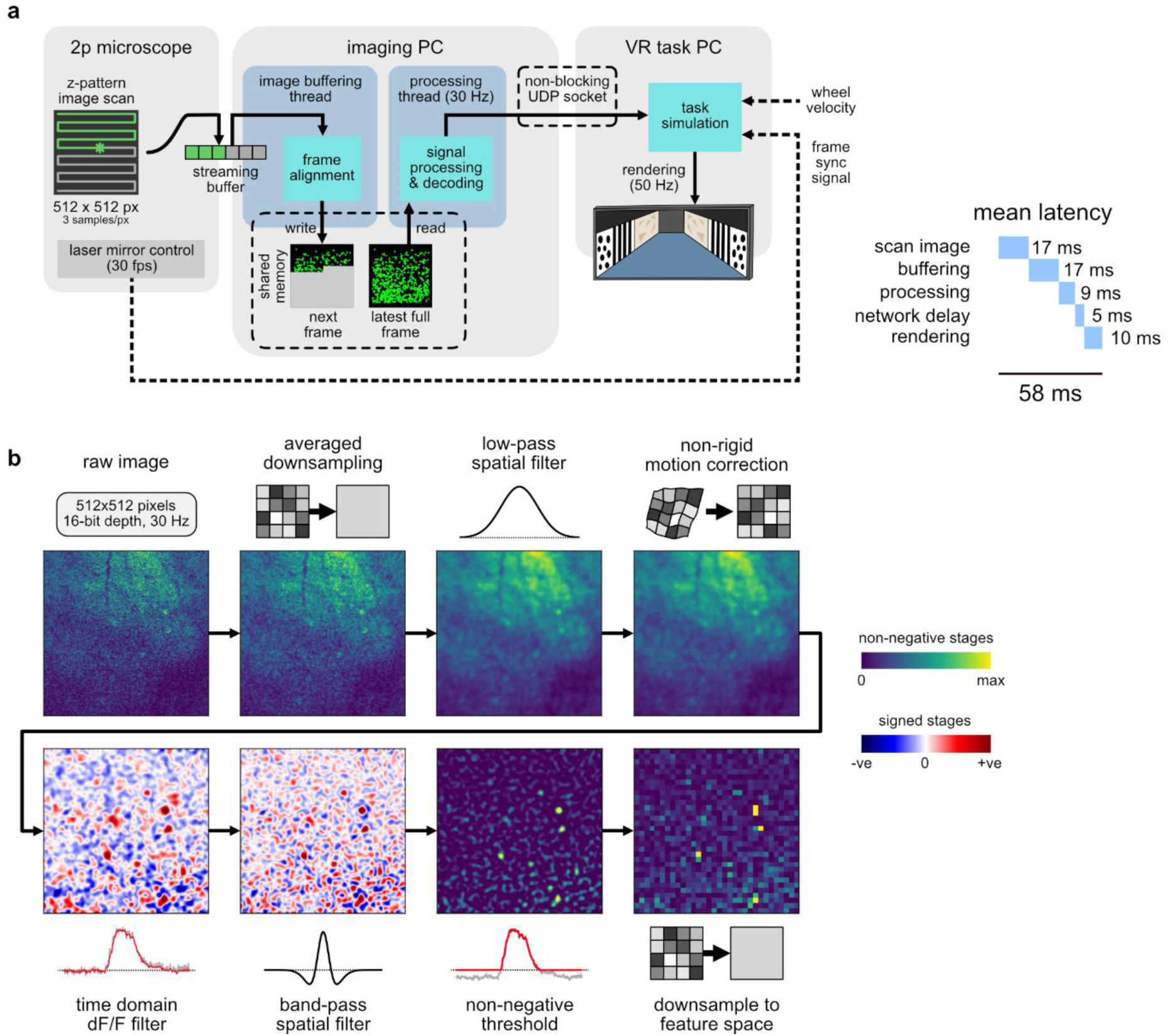
Schematic overview of the real-time BMI architecture. **(a)** A view of the role of each of the three major pieces of hardware (microscope, dedicated imaging PC, and PC dedicated to simulation of the VR environment) and which software components of the BMI pipeline they ran. Mean latency for each stage of the pipeline is shown estimated from the moment a neuron fires to its potential consequences are displayed on the screen. A side-channel directly from the microscope to the VR task PC ensured imaging frames could be synchronized to VR task variables. **(b)** Detail of the signal processing pipeline, illustrating the successive transformations used to extract the feature space applied to an example imaging frame.

**Supplementary Fig. 2.**
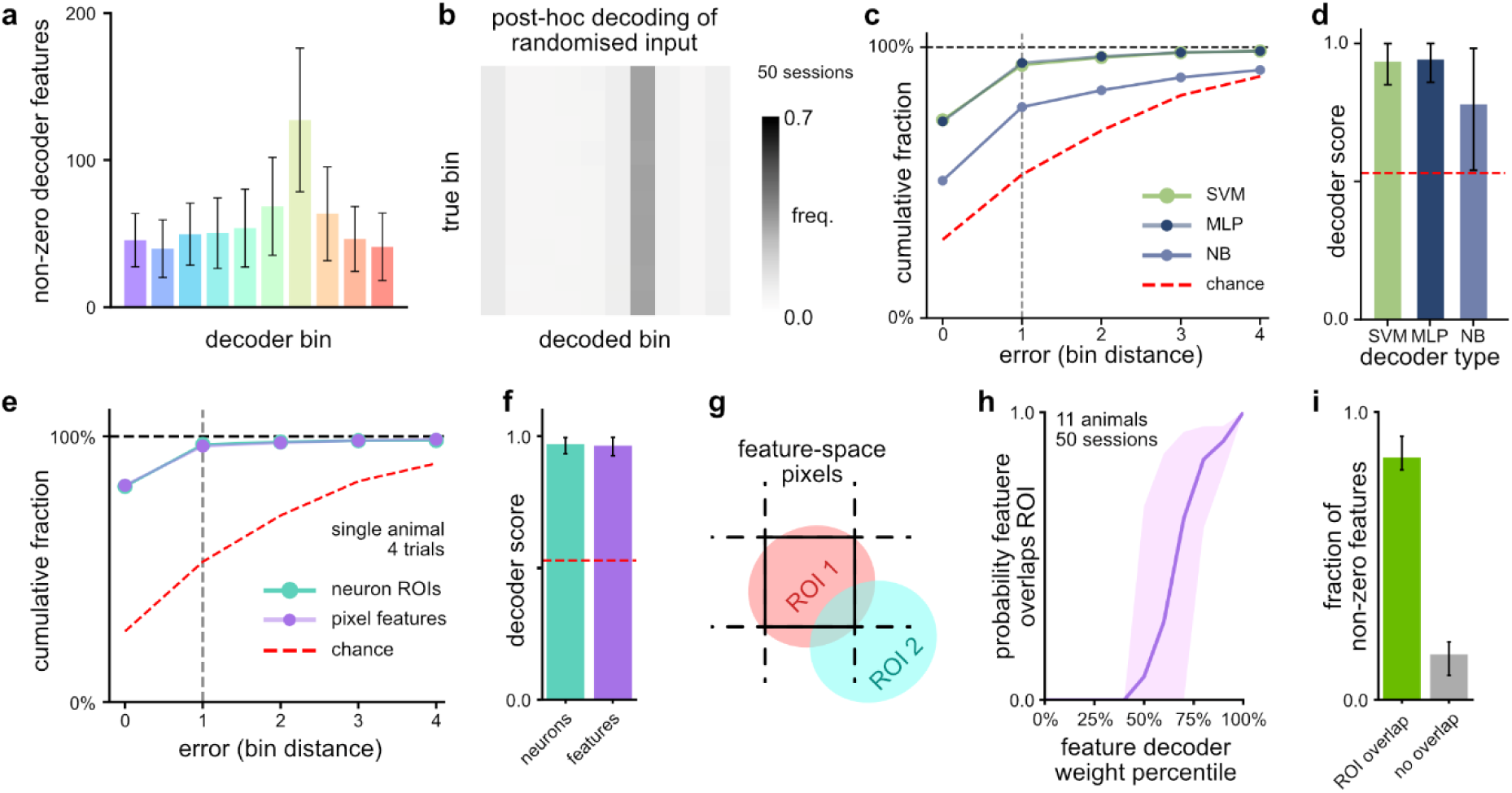
Offline evaluations of BMI decoders highlight performance characteristics of SVM classification. **(a)** The mean number of decoder features with non-zero weights for each spatial bin, shown over the 50 decoders trained on block 1 in locomotion + BMI sessions. Error bars indicate +/- SEM. Confusion matrix produced post-hoc by replacing the feature space with independent Gaussian noise and then evaluating decoder performance in block 2. Aggregated over all 50 locomotion + BMI sessions. **(c)** Decoding accuracy evaluated offline, trained and evaluated within 8 locomotion blocks (tested on unseen laps from those blocks), quantified by measuring the distance (in spatial bins) between predicted and true bin. Shown for three different classifiers: a support vector machine (SVM), a feedforward neural network (MLP), and a Naïve Bayes classifier (NB). Chance illustrated by a dummy decoder that predicts the most frequent bin. **(d)** The mean simplified decoder scores for (c), with error bars indicating min/max performance over the 8 blocks. **(e)** A comparison, as in (c), illustrating the equivalence between decoding from the signal processing feature space based on pixels, and directly from neuron ROIs extracted post-hoc with suite2p. Shown for 4 locomotion sessions from a single animal. **(f)** As in (d), shown for the comparison in (e). **(g)** A cartoon illustrating that the alignment between neuron ROIs and the pixel features that result from the signal processing pipeline are not guaranteed to line up perfectly, and that multiple neuron ROIs can overlap and therefore affect the same feature pixel. **(h)** The fraction of pixel features that overlap an underlying ROI, shown by ascending decoder weight percentile within that session. Line shows mean over 50 sessions, shaded region shows 5^th^ to 95^th^ percentiles over sessions. Decoders shown were trained on block 2 of locomotion + BMI sessions. **(i)** For the same decoders as (h), the ratio of features that do and do not overlap a neuron ROI for features with a non-zero decoder weight. Error bars show +/- SEM.

**Supplementary Fig. 3.**
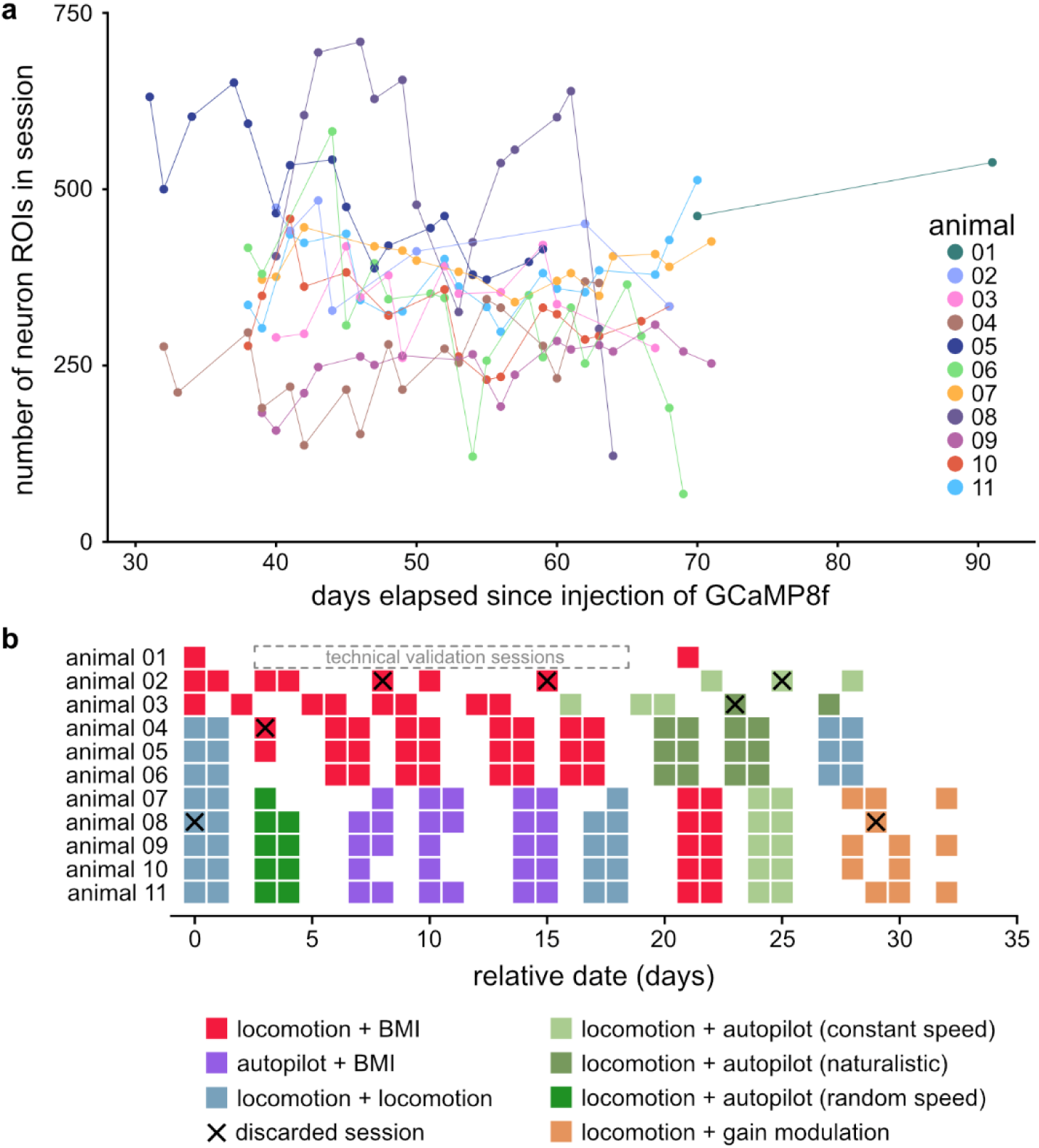
Overview of imaging schedule and quality between animals. **(a)** The number of neuron ROIs identified with suite2p in each experimental session for each animal. **(b)** Experimental schedule of each animal, showing session type relative to the first day shown in (a). Animal 01 was used for a series of technical validations to ensure correct functioning of the BMI system both prior to and between the two locomotion + BMI sessions shown in this panel. As experience in the VR environment can influence hippocampal representations, we explicitly show sessions that were discarded or aborted for technical reasons. Animal 02 had two sessions excluded from analysis due to an insufficient number of laps run in block 4 (but these sessions remain available in our dataset) and one session with image registration issues. Animal 03 had one session discarded due imaging issues. Animal 04 had one session discarded due to uncertainty about frame alignment between VR environment and microscopy data. Animal 08 had imaging registration issues in its first session, and its final imaging session was discarded due to fall-off in indicator response.

**Supplementary Fig. 4.**
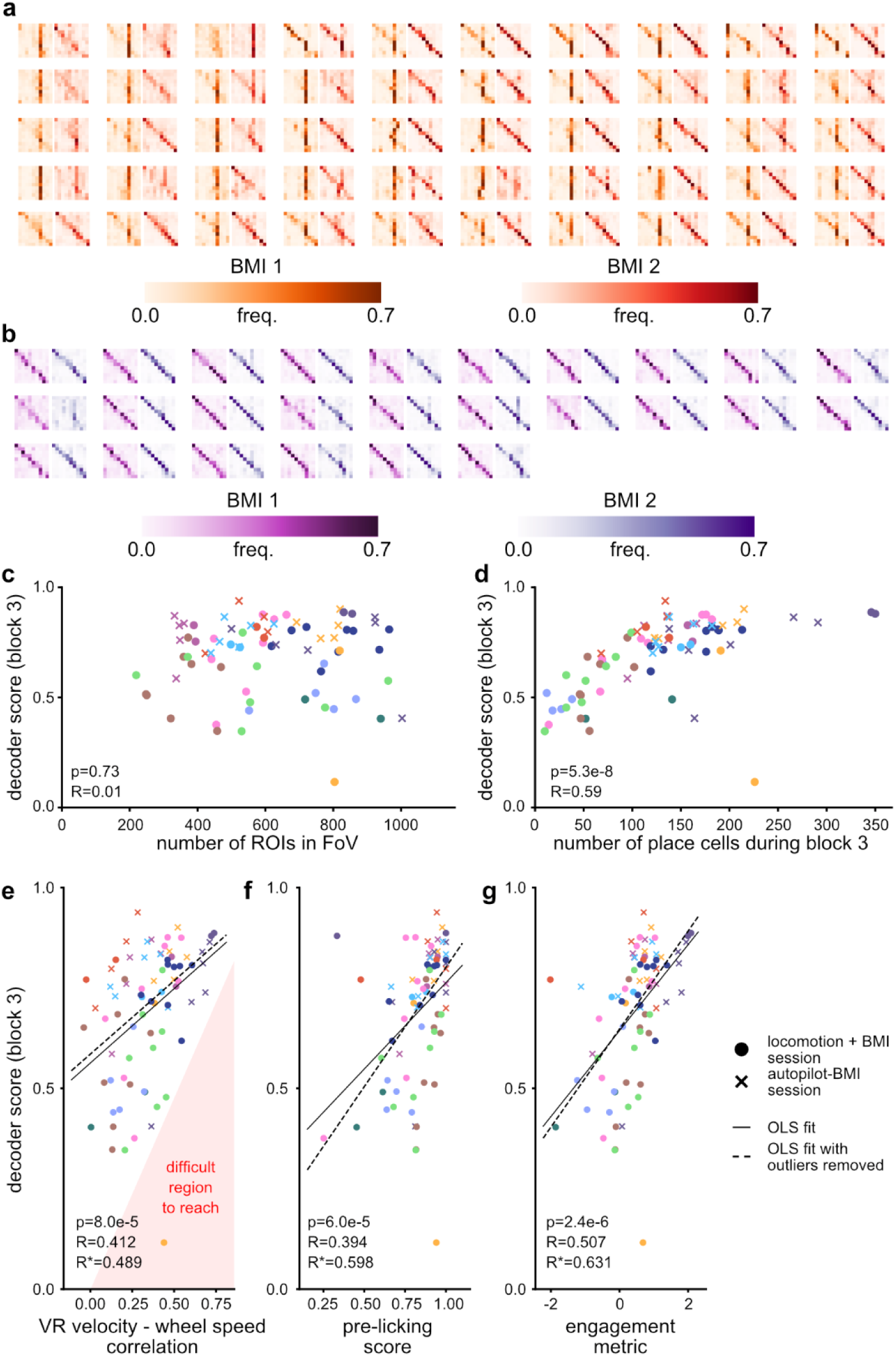
Individual variance in closed-loop BMI performance can be attributed to place cell count and task engagement. **(a)** Confusion matrices, as in Fig. 2e-f, shown for each locomotion + BMI session and **(b)** each autopilot + BMI session. Closed-loop decoder scores are shown for block 3 of each session against the number of ROIs visible in the FoV and **(d)** the number of cells with place fields during block 3. Circular markers indicate locomotion + BMI, crosses indicate autopilot + BMI. These same scores are also shown plotted against **(e)** the correlation between how the animal manipulated the wheel and the speed through the environment (noting that there is a region of this plot that is difficult to reach, as it requires the animal to match its running velocity to an unpredictable decoder), **(f)** the licking score within that block, and **(g)** an engagement metric which combines the variables from (e) and (f). R-values are reported for an ordinary least squares fit over all samples (solid lines), in addition to a fit to a subset removing outlier samples (dashed lines, R-values indicated as R*).

**Supplementary Fig. 5.**
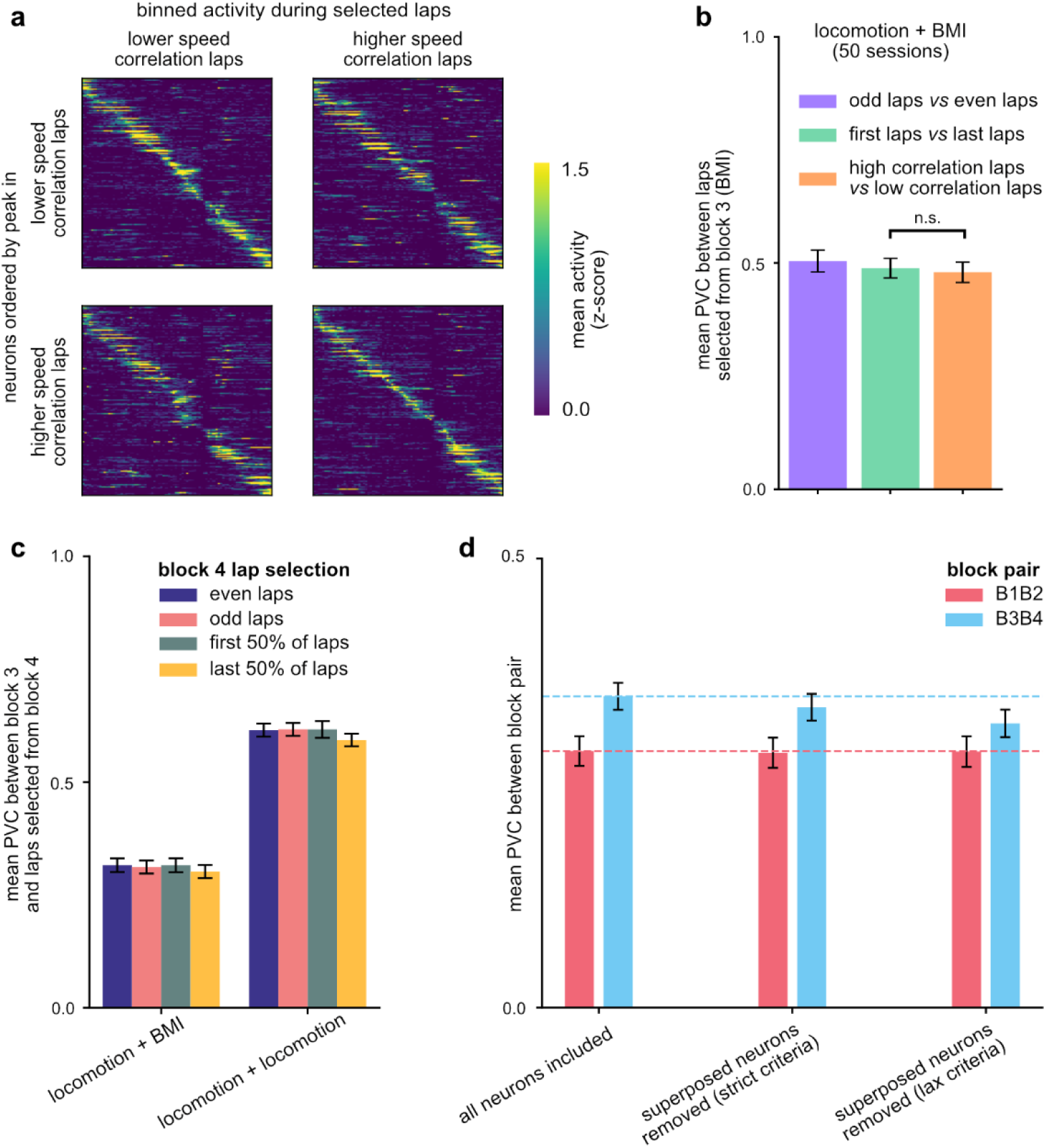
Validation of population-level changes in tuning during and following BMI usage. **(a)** Population ratemaps from within block 3 (BMI) of the example session shown in Fig. 2b, constructed by splitting the block into laps. The laps were ranked by the correlation between wheel speed and VR speed, and subsequently partitioned into the top half and the lower half of the ranking. **(b)** Applying the lap-splitting method from (a) to all 50 locomotion + BMI sessions (orange), and comparing it to splitting laps instead from the first and second half of the block (green) or odd and even laps (purple) by evaluating the mean PVC between the resultant ratemaps. Error bars show +/- SEM. There was no significant difference (p=0.10, uncorrected 2-sided paired t-test) between the mean PVC measured between the first and last laps of the block and the PVC measured between the high and low speed correlation laps. **(c)** A comparison, for both locomotion + BMI (50 sessions) and locomotion + locomotion (30 sessions), of the PVC between the ratemap from block 3 and ratemaps constructed from four different subsets of block 4: even laps, odd laps, the first half of laps, and the second half of laps. This illustrates in both conditions a small turnover of representation with time. Error bars show +/- SEM. **(d)** The mean PVCs between blocks 1 and 2 and between blocks 3 and 4, as previously described for Fig. 3a, showing how these change when neurons heuristically categorized as exhibiting superposition are withheld from the computation. Error bars show +/- SEM over the 50 sessions.

**Supplementary Fig. 6.**
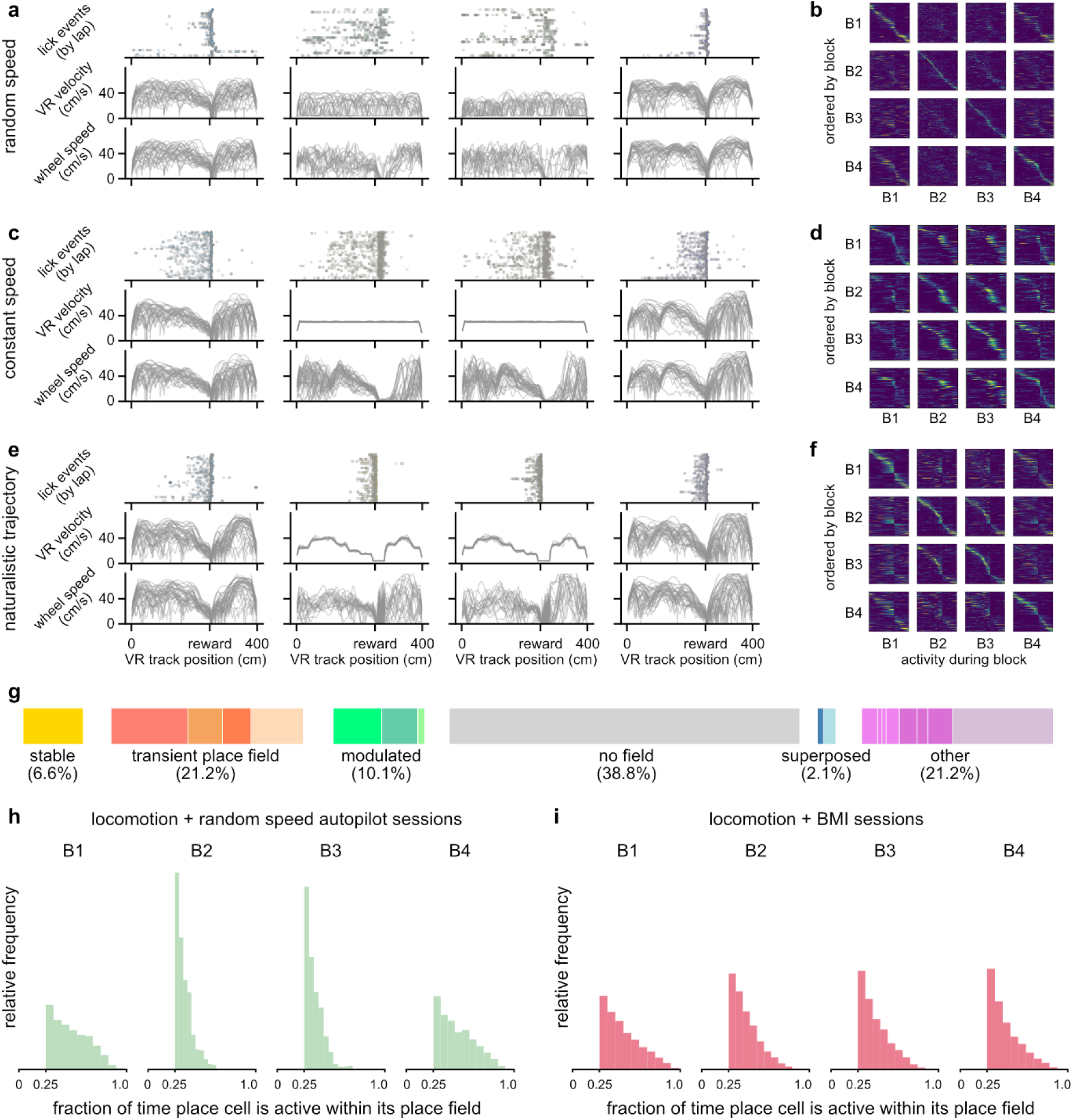
Details of the autopilot control conditions. **(a)** Traces of animal behavior during a single example locomotion + random autopilot session, showing lapwise spout-licking events and traces of both VR velocity and wheel manipulation speed. **(b)** A population ratemap for an example locomotion + random autopilot session. **(c)** As in (a), for a locomotion + constant speed autopilot session. **(d)** As in (b), for a locomotion + constant speed autopilot session. **(e)** As in (a), for a locomotion + naturalistic trajectory session. **(f)** As in (b), for a locomotion + naturalistic trajectory session. **(g)** The result of applying the same heuristic classification rules as in Fig. 3d to all locomotion + autopilot sessions (13160 neurons, 39 sessions). **(h)** Histograms showing, within each block of sessions using the random speed autopilot as the probe condition, the distributions of place field ‘quality’, as measured by the fraction of time the cell is active when the animal is within the corresponding place field. There is a discontinuity at 0.25, as neurons not meeting this threshold are rejected as candidate place cells. **(i)** As in (h), shown for the locomotion + BMI sessions.

**Supplementary Fig. 7.**
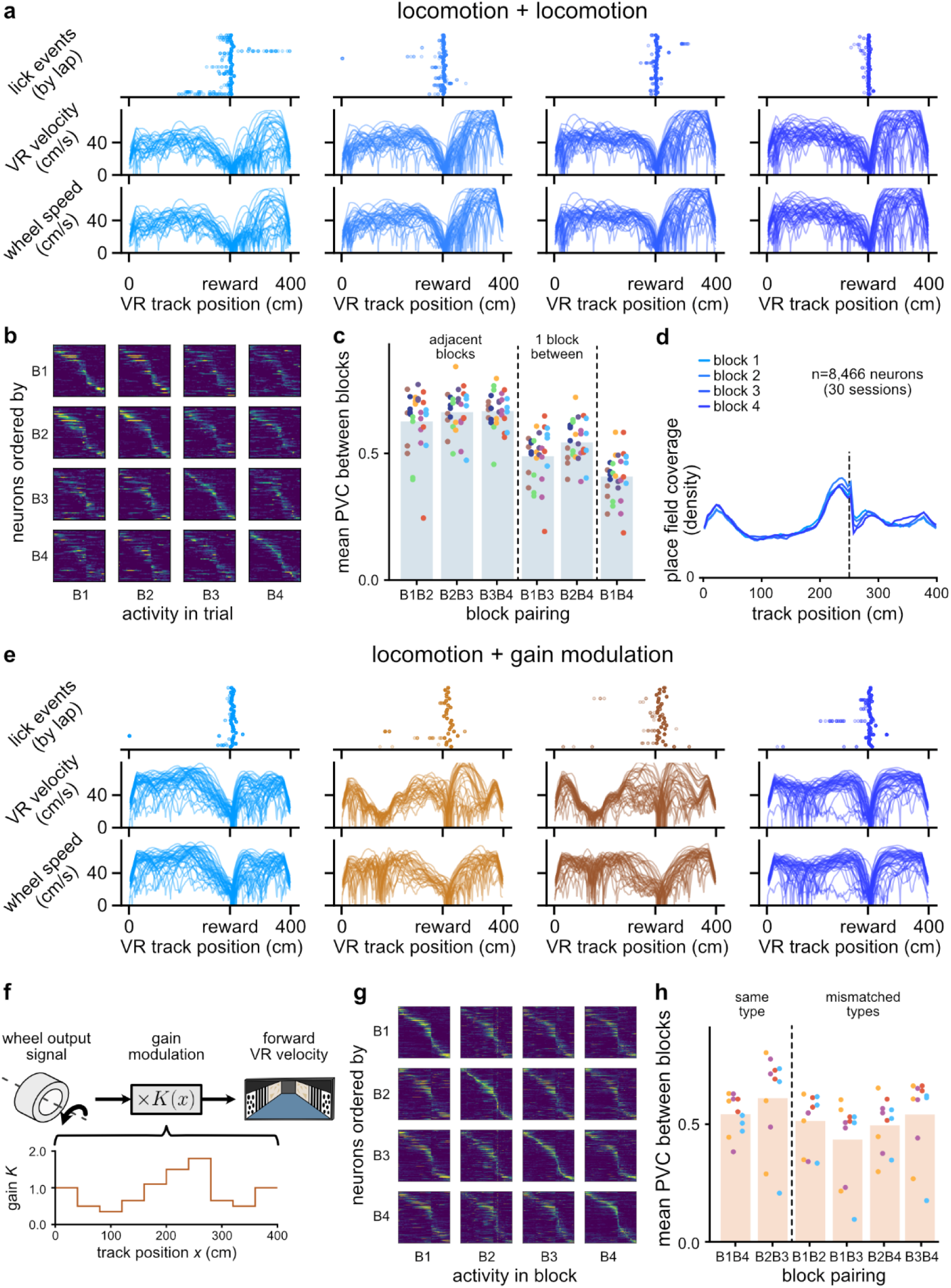
Additional controls in which animals retain access to agency and locomotion signals throughout the session. **(a)** Behavior traces, as in Fig. S6a, for an example locomotion + locomotion session. **(b)** A population ratemap for an example locomotion + locomotion session. **(c)** Mean population vector correlations between each block pairing, as in Fig. 3a and 4h, for all 30 locomotion + locomotion sessions (scatter point color indicates animal identity). Columns have been organized to group together block pairs that are 1, 2, and 3 blocks apart in time. **(d)** Relative coverage of the track by place fields, as in Fig. 4j, for the locomotion + locomotion session types. **(e)** As in (a), for a locomotion + gain modulation session. **(f)** Schematic illustrating how the gain modulation condition either amplifies or reduces the amount of VR traversal generated by wheel-running as a function of location on the track. **(g)** As in (b), for a locomotion + gain modulation session. **(h)** As in (c), for all locomotion + gain modulation sessions. This panel uses the column ordering convention from Fig. 3a and Fig. 4h.

**Supplementary Fig. 8.**
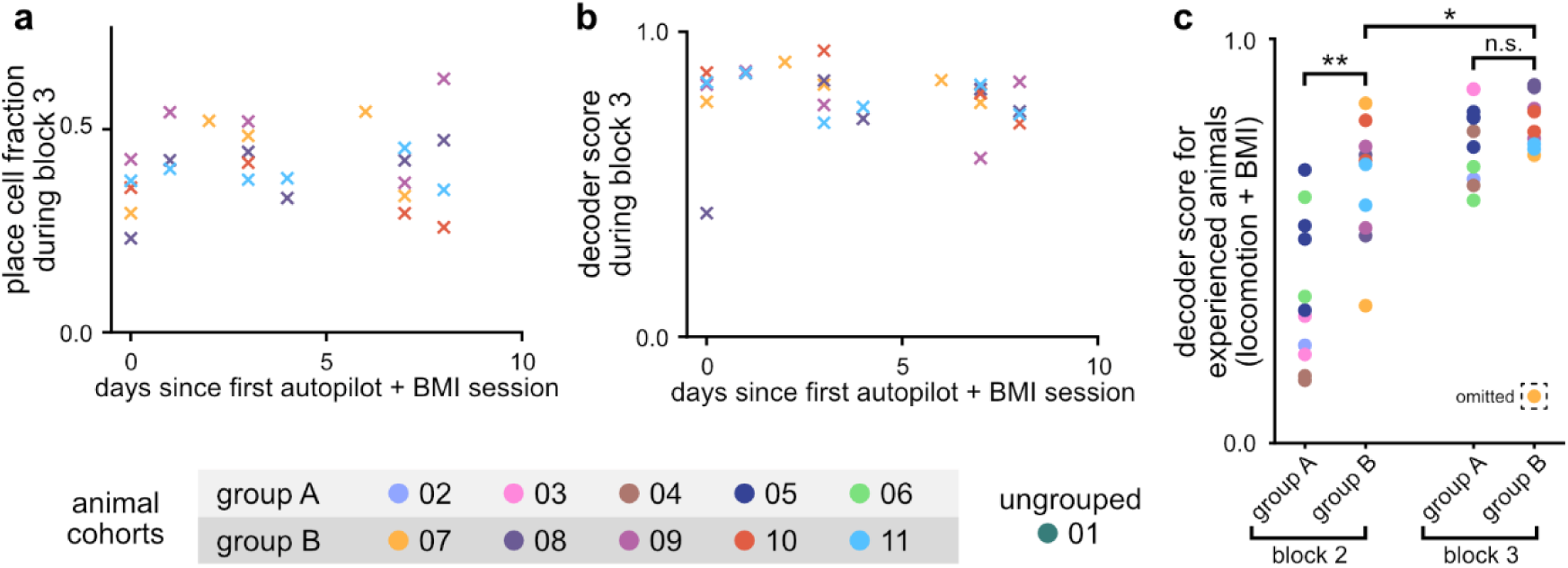
Animals with repeated exposure to the autopilot + BMI condition do not improve decoder accuracy with experience. **(a)** Fraction of the population that form place cells and **(b)** decoder score shown against increasing amounts of experience in the BMI condition, as shown in Fig. 2i, for the cohort of animals experiencing autopilot + BMI sessions. **(c)** A comparison between the two cohorts, shown after group B moved to locomotion + BMI sessions, of BMI decoder performance both during block 2 (pre-retraining) and block 3 (post-retraining). Shown for experienced animals (at least 10 days since first BMI experience) only. Despite group B demonstrating higher decoder scores than group A in block 2 (p=3.6e-4, two-sided independent t-test), and despite group B also experiencing an improvement in score between blocks 2 and 3 (p=0.02, same test), groups A and B did not differ significantly in performance in block 3 (p=0.30, same test).

**Supplementary Table 1:** A summary of results of statistical testing applied to the mean PVCs evaluated between different block pairings within several session types. Test 1 assesses whether there is a significant difference when comparing block pairings of the same condition (e.g. locomotion-locomotion or BMI-BMI) with pairings that mix conditions (e.g. locomotion-BMI). Test 2 assesses whether there is a significant difference in mean PVC when comparing the two pairings that match condition. Test 3, which we use as a way of assessing whether experience of the probe condition alters the baseline condition, tests whether the similarity between the initial baseline condition paired with the initial probe condition differs from the final probe paired with the final baseline condition (i.e. a ‘before and after’ check). The ‘locomotion + any autopilot’ row repeats the tests across merging autopilot conditions to determine if test 3 is still n.s. with larger N. For test 3, p-values are also shown without the Bonferroni correction.

| Session type | Test 1<br>Same vs different<br>control modality | Test 2<br>B1B4 ≠ B2B3 | Test 3<br>B1B2 ≠ B3B4 |
| --- | --- | --- | --- |
| locomotion + BMI | p=7.35e-5 | p=0.46 (n.s.) | p=0.020<br>(p=3.37e-3 uncorrected) |
| locomotion + constant autopilot | p=4.34e-4 | p=1.21e-3 | p=1.0 (n.s.)<br>(0.59 uncorrected) |
| locomotion + naturalistic autopilot | p=0.012 | p=2.46e-3 | p=0.48 (n.s.)<br>(0.08 uncorrected) |
| locomotion + random autopilot | p=6.57e-4 | p=0.028 | p=1.0 (n.s.)<br>(0.42 uncorrected) |
| locomotion + any autopilot | p=6.60e-8 | p=1.58e-3 | p=1.0 (n.s.)<br>(0.47 uncorrected) |
| random autopilot + BMI | p=3.58e-5 | p=2.56e-5 | p=1.0 (n.s.)<br>(0.28 uncorrected) |

**Supplementary Movie 1. Three example consecutive laps from block 3 of a locomotion + BMI session, illustrating BMI performance of the navigation task.** On the first lap, the mouse slightly overshoots the reward site. On the second lap, it moves the wheel in close agreement with how the BMI drives motion through the environment. On the third lap, it stops moving on the wheel during a portion of the traversal while BMI-driven motion continues at high speed. From left to right, the top row shows a video recording of the animal, the raw population imagery, and the feature space used as input to the SVM decoder in the BMI. The bottom row shows traces of the animal’s VR traversal speed as driven by BMI (included a black dashed line to indicate the target trajectory) and traces of the animal’s running speed on the wheel. The final plot shows the animal’s location on the track as a function of time, with the track divided into the 10 spatial bins used for decoding. The color of each point indicates the output class of the BMI decoder at each timestep.

